# Anti-diabetic drug Repaglinide induces Apoptosis, Cell Cycle Arrest, and Inhibits Cell Migration in Human Breast and Lung Cancer Cells

**DOI:** 10.64898/2026.02.25.707939

**Authors:** P K Hima, K Aswathy, Nagendra Yarla, Govinda Rao Duddukuri

## Abstract

**Introduction:** Drug repurposing offers a cost-effective and time-efficient strategy for cancer therapy by leveraging existing drugs with established safety profiles, thus functioning as an alternative therapeutic strategy in demanding diseases such as cancer. Antidiabetic agents, in particular, have demonstrated encouraging anticancer potential. Among them, the non-sulfonylurea insulin secretagogue repaglinide (RPG) has shown emerging anticancer potential, yet its effects on breast and lung cancers remain largely unexplored. Thus, this study investigates the anticancer activity of repaglinide in human breast (MCF-7) and lung (A549) cancer cell lines, focusing on its cytotoxic, pro-apoptotic, anti-proliferative, and anti-migratory effects and the underlying possible molecular mechanisms.

**Methodology and Results:** MTT cytotoxic assay revealed that RPG reduced cell viability in a dose-/time-dependent manner, with an IC₅₀ (48h) of 100.8 ± 3.98 µM for MCF-7 and 104 ± 3 µM for A549. Further, the apoptotic effect of RPG on both cell lines was evidenced by double staining assays, comet assay, and western blotting analysis, suggesting that RPG explicitly caused DNA damage and activated intrinsic and extrinsic apoptosis pathways. Additionally, RPG suppressed clonogenicity and enforced G1 arrest in MCF7 and A549 cells by modulating cell cycle regulations as well as cell proliferation pathways. Moreover, RPG markedly suppressed cell motility, as demonstrated by scratch and Transwell migration/invasion assays, which is correlated with reduced MMP-2 and MMP-9 expression, confirmed by gelatin zymography and western blotting.

**Conclusion:** Conclusively, Repaglinide exerts potent anticancer effects in breast and lung cancer cells by modulating key oncogenic signaling pathways, and thus can be considered a promising candidate for repurposing in cancer therapy.

## 1. Introduction

Cancer remains a major global health burden and one of the leading causes of morbidity and mortality worldwide, characterized by uncontrolled cell proliferation, genomic instability, metabolic reprogramming, and complex molecular heterogeneity [1,2]. Although advances in surgery, chemotherapy, radiotherapy, and targeted therapies have improved patient management, many cancers continue to exhibit therapeutic resistance, recurrence, off-target toxicity, and substantial treatment costs [3,4]. These persistent challenges underscore the need for innovative, mechanism-based, and economically feasible therapeutic strategies.

In this context, drug repurposing has emerged as a rational and strategically advantageous approach in oncology. By identifying novel anticancer applications for already approved drugs, this strategy reduces the time, cost, and high failure rates associated with conventional drug development while leveraging established safety and pharmacokinetic profiles [5,6]. Drug repurposing in oncology is particularly compelling when supported by robust epidemiological associations and shared molecular mechanisms between the drug’s primary therapeutic indication and tumorigenesis. In this regard, a substantial body of evidence has established a significant link between type 2 diabetes mellitus (T2DM) and cancer development and progression [7]. Epidemiological evidence indicates that T2DM is associated with an elevated risk of several malignancies, with particularly strong associations observed in breast and lung cancer. Breast carcinoma remains the foremost cause of cancer-related mortality among women globally, and women with T2DM demonstrate approximately a 23% greater likelihood of developing breast cnacer [8,9]. Likewise, individuals with T2DM exhibit a 20–48% elevated relative risk of lung cancer [10]. These clinical observations are reinforced by shared biological mechanisms that mechanistically connect metabolic dysfunction to tumorigenesis. Chronic hyperglycemia, hyperinsulinemia, and insulin/IGF-1–mediated mitogenic signaling activate oncogenic cascades such as PI3K/Akt and K-Ras, promote oxidative stress and inflammation, and create a tumor-permissive microenvironment [7,11]. The integration of metabolic dysregulation with proliferative and survival signaling provides a strong mechanistic basis for evaluating metabolic modulators as anticancer agents.

Consistent with this rationale, several classes of antidiabetic drugs, including biguanides (metformin), thiazolidinediones (pioglitazone), and SGLT2 inhibitors (canagliflozin), have demonstrated significant anticancer properties beyond glycaemic control. These agents have been reported to modulate cellular metabolism, suppress proliferative signaling pathways, induce apoptotic cell death, and inhibit inflammatory responses and angiogenesis [7,12–14]. In contrast to these well studied classes, insulin secretagogues, comprising sulfonylureas and meglitinides, represent a comparatively underexplored yet mechanistically relevant group in oncology. Sulfonylureas such as Glibenclamide and Gliclazide have demonstrated antiproliferative and pro-apoptotic effects in experimental cancer models, including breast and lung cancers, through induction of oxidative stress, mitochondrial dysfunction, and modulation of key survival pathways such as AKT and JNK [4,15]. These findings suggest that insulin secretagogues may exert direct tumor-suppressive actions independent of their systemic metabolic effects.

Extending this rationale, insulin secretagogues, particularly meglitinides, warrant further investigation in the oncologic setting. Meglitinides, including repaglinide (RPG), are non-sulfonylurea insulin secretagogues that target the sulfonylurea receptor 1 (SUR1) component of ATP-sensitive potassium (K^+^ATP) channels in pancreatic β-cells [16, 17]. Unlike extensively investigated antidiabetic classes, repaglinide remains relatively underexplored in cancer research despite several attributes that strengthen its translational potential. Clinically, RPG possesses a well-characterized safety profile, favorable pharmacokinetics, and widespread therapeutic use, thereby reducing developmental uncertainty and facilitating potential clinical application. Importantly, accumulating preclinical data demonstrate that repaglinide exerts direct antitumor effects across multiple malignancies. In glioblastoma models, RPG suppresses cellular proliferation and triggers caspase-dependent apoptotic pathways while attenuating pro-survival signaling cascades [18]. In hepatocellular carcinoma, RPG has been shown to inhibit tumor growth, reduce migratory potential, and promote apoptotic cell death, potentially via modulation of mitochondrial integrity and oxidative stress [19]. Additionally, emerging evidence in neuronal cancer models indicates that repaglinide interferes with cell cycle progression and metastatic behaviour [20] . The reproducibility of these antiproliferative and pro-apoptotic effects across biologically distinct tumor systems suggests that repaglinide may target core regulatory mechanisms fundamental to cancer progression. Collectively, these findings highlight the broad-spectrum anticancer activity of Repaglinide across diverse tumor types, demonstrating its ability to regulate key pathways involved in cellular proliferation, survival, and metastatic progression. However, despite growing evidence of its anticancer activity in multiple tumor models, the therapeutic efficacy and underlying molecular mechanisms of RPG in breast and lung cancers remain insufficiently defined. Given RPG demonstrated multi-targeted antitumor effects and favourable clinical profile, a systematic evaluation of repaglinide in these malignancies is therefore both timely and scientifically justified. Therefore, the present study aims to investigate the cytotoxic, antiproliferative, and anti-migratory effects of repaglinide in breast and lung cancer cell lines *in vitro,* thereby evaluating its potential repositioning as a mechanistically informed anticancer agent.

## 2. Materials and methods

### 2.1. Chemicals

MTT (3-(4,5-dimethylthiazol-2-yl)-2,5-diphenyltetrazolium bromide), purchased from Himedia Pvt., Ltd., Mumbai, India. Dead Cell Apoptosis Kits with Annexin V for Flow Cytometry (Cat No# V13242) and Propidium iodide, Invitrogen (Carlsbad, CA, USA). Acridine orange and Crystal violet were purchased from Sigma-Aldrich, and Ethidium bromide was purchased from Bio-Rad. The details of the antibodies used for the study are listed in the Supplementary Table 1. RPG Mother stock preparation: Repaglinide (RPG), Cat no# R9028-50MG, was purchased from Sigma-Aldrich (St. Louis, MO, USA) and dissolved in dimethyl sulfoxide (DMSO). The overall DMSO concentration does not exceed 0.1% in all the experiments. All experiments were conducted using sublethal doses (below IC₅₀), determined based on observed morphological changes. Pifithrin-α treatment: Pifithrin-α (PFR-α), Cat no# **63208-82-2** (5 mg), was purchased from Mechem Express (MEC) and dissolved in DMSO. A concentration of 10 µM pifithrin-α was selected for co-treatment, in accordance with dosages commonly employed in prior studies [21,22]

### 2.2. Cell culture

Human breast cancer (MCF-7) and lung cancer (A549) cell lines were obtained from the National Center for Cancer Cell Lines (NCCS, Pune, India). The cells were cultured in DMEM (Dulbecco’s Modified Eagle Medium), with 10% FBS (fetal bovine serum), 2 µM L-glutamine, and 100 µg/mL antibiotic–antimycotic solution (HI Media Pvt. Ltd., Mumbai, India). The flasks were then maintained at 37 °C in a humidified incubator with 5% CO₂. Cells were routinely subcultured in 100 mm culture dishes and harvested by trypsinization for various experiments upon reaching approximately 85% confluence.

### 2.3. Cell Cytotoxicity Assay- MTT Assay

MCF-7 and A549 (5 × 10³ cells per well) were seeded in 96-well plates, with each condition performed in triplicate. After 20 hours of incubation, the cells were treated with varying concentrations of repaglinide (RPG), starting from 500 µM, with control wells receiving 0.1% DMSO and further incubated for 24 and 48 hours. Following RPG incubation, cells were treated with MTT [3-(4,5-dimethylthiazol-2-yl)-2,5-diphenyltetrazolium bromide] reagent (5 mg/mL in PBS) and incubated for another 3 hours. After dissolving the formazan crystals in DMSO, the absorbance was measured at 570 nm, with 630 nm serving as a reference to correct for background noise. A graph was plotted with cell viability against RPG concentration to calculate the IC50 values [23]

### 2.4. Morphology assessment

To assess the effect of RPG on cellular morphology, MCF-7 and A549 cells were seeded in a 6-well plate, followed by treatment with various sublethal concentrations of RPG (0–80 µM) and incubated for 24 and 48 hours. Post-treatment, morphological changes were observed and documented using a Carl Zeiss inverted microscope (Carl Zeiss Microscopy GmbH, Jena, Germany) at 20X magnification, and representative images were captured for analysis. The concentration with significant morphological alteration was selected for further experiments.

### 2.5. Acridine Orange/ Ethidium Bromide (AO/EtBr) Dual Staining

The apoptotic morphology of RPG-treated cells was assessed using acridine orange/ethidium bromide (AO/EtBr) staining. MCF-7 and A549 cells were treated with two sublethal concentrations of RPG (40 µM and 80 µM), then incubated for 24 and 48 hours. Following treatment, cells were harvested, resuspended in ice-cold PBS, and stained with 20 µL of AO/EtBr staining solution (1:1v/v; AO-5mg/mL and EtBr-10mg/mL) for 20 minutes. 20 µL of the solution was then mounted with a coverslip on a clean glass slide. Stained cells were subsequently observed under a fluorescence microscope (Zeiss Axio Observer fluorescence microscope) using the blue channel to detect AO-stained (viable or early apoptotic) cells and the green channel for EtBr-stained (late apoptotic or necrotic) cells. ImageJ software quantified the number of apoptotic cells in both control and treated groups, and a graph was plotted.

### 2.6. Quantitative Analysis of Apoptosis using Annexin V-FITC/PI Staining

Apoptotic cell populations were quantitatively assessed by flow cytometry using the Annexin V-FITC/PI apoptosis detection kit (Invitrogen, Thermo Fisher Scientific, USA), as per the manufacturer’s instructions. Briefly, MCF-7 and A549 cells were seeded in 6-well plates and treated with or without RPG (40 μM and 80 μM). Following treatment, 48 hours of incubation, cells were collected and resuspended in 100 µL of 1× binding buffer, stained using 5 µL of Annexin V-FITC and 5 µL of propidium iodide (PI) for 20 minutes at 37 °C in the dark. Subsequently, 400 µL of 1× binding buffer was added, and the apoptotic populations were analyzed using flow cytometry (FACS #PARTEC CYFLOW, space software- FlowMax Sysmex Partec GmbH, Görlitz, Germany)

### 2.7. DNA damage assessment using Comet Assay

A neutral comet assay was conducted to evaluate the DNA-damaging effects of RPG in MCF-7 and A549 cells, as explained elsewhere [24]. Briefly, 3 × 10⁶ cells were plated in 6-well plates and treated with two sublethal concentrations of RPG for 48 hours. Post-incubation, the cells were harvested and resuspended in 200 µL of ice-cold PBS. The cell suspension was mixed with 1% agarose, spread onto pre-coated glass slides, and subsequently air-dried at room temperature for 20–30 minutes. The slides were then immersed in neutral lysis buffer (0.5 M Na₂EDTA, 2% SDS, 0.5 mg/mL proteinase K, pH >8) and incubated overnight in the dark. After lysis, slides were washed twice with 1× TAE buffer and subjected to electrophoresis in 1× TAE at 0.6 V/cm for 25 minutes. Subsequently, slides were stained with propidium iodide (PI) for 20 minutes and visualized under a fluorescence microscope at 20X magnification. Random pictures were taken from different sites, and a total of 50 comets were used for analysis. The comet length and other parameters were quantified using CASP-Lab (Comet Assay Software Project) software.

### 2.8. Colony formation assay

The effect of RPG on colony-forming ability was assessed using a clonogenic assay [25]. Cells (MCF7 and A549) were cultured in 6-well plates at a seeding density of 1.5 × 10³ cells/well and incubated overnight. Following the incubation, the cells were treated with either vehicle control or varying concentrations of RPG (0–100 µM) for 7 days, with media changed every 2 days. Following the incubation period, the media was aspirated, and the cells were fixed using 4% paraformaldehyde for 15 to 20 minutes and stained with 0.01% (v/v) crystal violet for 30 minutes. After staining, the wells were gently washed twice with distilled water and imaged. A stereomicroscope was used to count colonies containing ≥50 cells, and the surviving fraction was determined using the following formula S.F = PE of treated samples/ PE of control PE (plating efficiency) = Number of colonies counted/ number of cells

### 2.9. Cell cycle analysis using Flow cytometry

Flow cytometry based on DNA content analyzed the cell cycle distribution in response to RPG treatment. MCF-7 and A549 cells were seeded in 10 cm culture dishes at a density of 2.2 × 10⁶ cells per plate and treated with or without RPG for 24 hours. Following the incubation, cells were harvested by trypsinization, fixed in 70% ethanol by gentle vortexing, followed by storage on ice for a minimum of 2 hours. The cells were washed thrice with PBS, incubated with RNase A (10 mg/mL) at 37 °C for 1 hour. Afterward, they were stained with propidium iodide (PI, 10 mg/mL) for 20-30 minutes at room temperature in the dark. DNA content was analyzed using a CytoFLEX (Beckman Coulter, Brea, California, United States) flow cytometer and CytExpert software.

### 2.10. Scratch wound healing assay

MCF-7 and A549 cells were plated in 24-well plates in triplicate at a density of 2 × 10⁵ cells per well and allowed to form a uniform monolayer. Once confluence was reached, a scratch wound was created at the center of each well using a 10 µL micropipette tip. The wells were washed twice with PBS to remove detached cells and debris. Cells were treated with various concentrations of RPG (0–80 µM) in serum-free media, with vehicle-treated wells serving as controls. Following treatment, plates were incubated for 24 and 48 hours. Wound closure was monitored and imaged at 10X magnification. The wound area was quantified by drawing a reference line along the wound edges and measuring the remaining gap using ImageJ software.

### 2.11. Transwell migration and invasion assay

The effect of RPG on cellular migration and invasion was assessed *in vitro* using a transwell chamber assay, as outlined in previous studies [26]. Transwell chamber inserts with 0.8 µm pore size polycarbonate membranes (HiMedia Pvt. Ltd.) were used for the experiment. To perform the invasion assay, the upper chamber was pre-coated with 50 µL of Matrigel (Gibco Geltrex, Catalog No: A15696-01) and incubated at 37 °C for 1 hour prior to cell seeding. Further, MCF7 and A549 cells were harvested, resuspended in serum-free medium, and 100 µL of the cell suspension (2 × 10⁴ cells/well) was seeded into each chamber. The lower chamber was supplemented with 10% FBS-containing medium to act as a chemoattractant. After a 1-hour initial incubation, the cells were treated with two sublethal concentrations of RPG and further incubated for 24 hours. After incubation, the cells that had migrated or invaded to the bottom of the chambers were fixed using 4% paraformaldehyde and stained with 0.01% (v/v) crystal violet. The cells were then visualized, photographed, and counted using a stereomicroscope.

### 2.12. Gelatin Zymography

Gelatin zymography was performed to understand the effect of RPG on the enzymatic activity of MMP-2 and MMP-9 [26]. Briefly, MCF-7 and A549 cells were seeded in 6-well plates and treated with or without varying concentrations of RPG (20, 40, and 80 µM) for 48 hours. Following incubation, the culture supernatants were obtained and centrifuged to remove cellular debris. Protein quantification in the supernatants was determined using the Bradford assay [27]. 5 µg of total protein was loaded onto an 8% SDS-PAGE gel containing 0.1% gelatin, and electrophoresis was carried out under non-reducing conditions at room temperature for 2 hours. Subsequently, the gel was incubated in renaturing buffer (2.5% Triton X-100) for 30 minutes at room temperature with gentle agitation to restore enzymatic activity. The gel was then washed and incubated in developing buffer (50 mM Tris base, 0.2 M NaCl, 5 mM CaCl₂, 0.02% Brij 35, 1 µM ZnCl₂, pH 7.6) for 16 hours at 37 °C. Post incubation, the gel was stained with Coomassie Brilliant Blue R for 30-40 minutes and destained with distilled water. Proteolytic activity appeared as clear bands against a blue background. The gel images were captured, and the band intensities were measured using ImageJ software. A parallel SDS-PAGE gel without gelatin was run under identical conditions using the same protein samples as a loading control.

### 2.13. Western blotting

After 48 hours of treatment with three sublethal concentrations of RPG (40, 60, and 80 µM), MCF-7 and A549 cells were harvested and lysed in RIPA lysis buffer supplemented with a protease inhibitor cocktail. Further, the total protein concentration in the collected samples was determined using the Bradford assay [27]. Equal amounts of protein (20 µg) were separated by SDS-PAGE (8-12%) and transferred onto a methanol-activated PVDF membrane (Bio-Rad Laboratories, USA) using a western blot transfer system (Bio-Rad, USA). Afterward, the membrane was blocked with 5% BSA and then incubated overnight (16 hours) at 4 °C with the appropriate primary antibodies with gentle agitation. Following primary incubation, the membrane was washed thrice with TBST and then incubated with appropriate HRP-conjugated secondary antibodies (anti-mouse or anti-rabbit, CST) for 1 hour, 30 minutes at room temperature with gentle shaking. After washing the membrane three more times with TBST, signals were developed using Clarity Western ECL substrate (Bio-Rad, Catalog No: 170-5060) and visualized with a C-DiGit blot scanner. Band intensities were quantified for densitometric analysis using GAPDH as an internal control.

### 2.14. Immunofluorescence studies

MCF-7 and A549 cells were seeded at a density of 0.4 ×10^6^ cells onto cell culture-grade coverslips and treated with 80 µM RPG, followed by incubation for 48 hours. After treatment, coverslips were removed and washed 2-3 times with TSB, then fixed with 4% paraformaldehyde for 20 minutes. Following the fixation, the cells were permeabilized with 0.1% Triton X-100 for 10 minutes. Subsequently, the blocking step was performed with 5% BSA at room temperature for 1 hour. The coverslips were washed with TSB and incubated with the appropriate primary antibodies overnight (16 hours) at 4 °C. After the incubation, the cells were rinsed thrice with TSB and incubated with Alexa Fluor 488 and/or 594 conjugated secondary antibodies for 1 hour in the dark. Nuclei were counterstained with DAPI (Sigma Aldrich: cat# D9542) for 30 minutes in the dark. The coverslips were then washed, mounted onto clean glass slides, and analyzed under the fluorescence microscope using oil immersion at 100× magnification.

### 2.15. Statistical analysis

Data were represented as mean ± SD of at least three independent experiments. Statistical analysis was performed with two-way ANOVA followed by Tukey’s Post-Test multiple comparisons using GraphPad PRISM (Version 10.0) software. The statistical significance is represented as *P<0.05, **P<0.01, ***P<0.001, and P***< 0.0001 compared with the respective control.

## 3. RESULTS

### 3.1. Repaglinide induces cell cytotoxicity and morphological changes in human breast and lung cancer cells

The anti-cancer effect of Repaglinide (RPG) was evaluated employing breast (MCF-7) and lung (A549) cancer cells. The cytotoxicity of RPG on both these cells was tested at 24 and 48 hours using the MTT colorimetric assay. As represented in Fig. 1A and 1B, RPG significantly reduces the viability of MCF-7 and A549 cells in a dose and time-dependent manner, with a more pronounced effect observed after 48h of treatment. The half-maximal inhibitory concentration (IC₅₀) values decreased with time, from 128 ± 1.9 to 100.8 ± 3.98 µM for MCF7, and from 143 ± 2.8 to 104 ± 3 µM for A549 cells, indicating enhanced sensitivity with prolonged exposure to RPG. Notably, both cell lines exhibited comparable sensitivity towards RPG treatment. These results signify the cytotoxic effect of RPG on both MCF7 and A549 cells.

**Figure 1:**
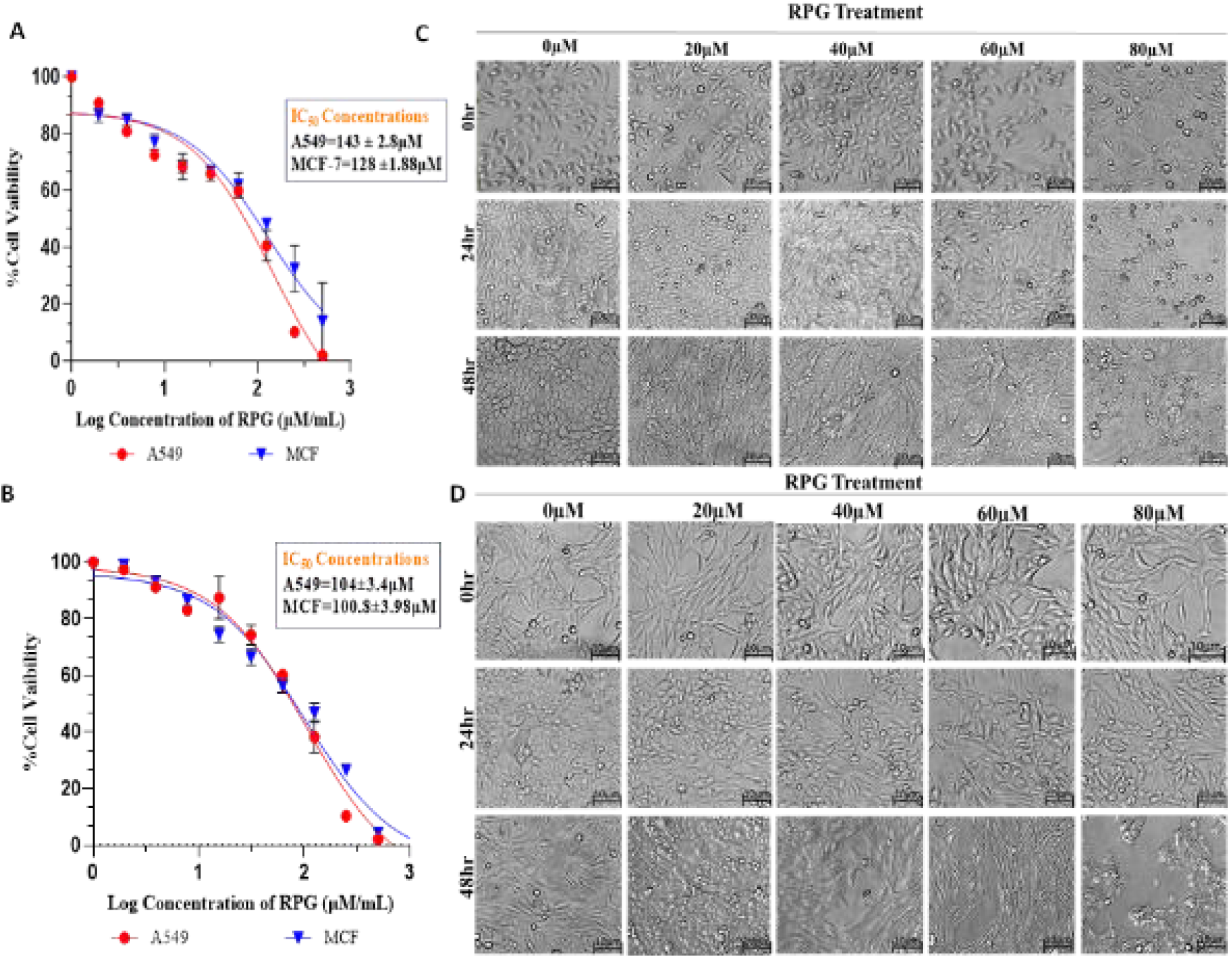
RPG induced cytotoxicity and morphological alterations in MCF7 and A549 cells. Cells were exposed to varying concentrations of repaglinide (RPG; 0–500 μM) for 24 and 48 hours, and cell viability was assessed using the MTT assay. The graph illustrates the cytotoxicity profile of RPG in MCF7 and A549 cells for 24 h (A) and 48 h (B). Results are expressed as a percentage of control ± SD (n=3). Panels C and D: Representative images showing the morphological alteration induced by the RPG on MCF7 and A549 cells, respectively. The normal morphology of the cells was significantly distorted after 48 hours, with cells showing elongated shapes and boundaries representing that of loosely adhered cells.

Considering the pronounced cytotoxic effects induced by RPG treatment, morphological alterations were systematically evaluated following exposure to sublethal concentrations of RPG (0, 20, 40, 60, and 80 µM). RPG treatment led to dose- and time-dependent morphological alterations in both cell lines, particularly after 48 hours. Cells exhibited progressive loss of structural integrity with increasing concentrations. Cells exposed to RPG (≥40 µM) displayed spindle-like shapes, reduced adhesion, and irregular membranes. At higher concentrations, prominent features of apoptosis, including membrane blebbing and structural disintegration, were evident in both cell lines compared to vehicle-treated controls. (Fig. 1C, 1D).

### 3.2. RPG Facilitates Apoptosis in MCF7 and A549 Cancer Cells

In order to evaluate the pro-apoptotic effects of Repaglinide (RPG), MCF-7 and A549 cells were treated with two sub-lethal concentrations of RPG (40 µM and 80 µM) and incubated for 24 and 48 hours. Apoptotic morphological changes were assessed using acridine orange/ethidium bromide (AO/EtBr) double staining. As depicted in Fig. 2A and 2B, RPG-treated cells displayed hallmark features of apoptosis in a dose- and time-dependent manner. Specifically, a substantial reduction in green-fluorescing viable cells was observed in RPG-treated samples compared to vesicle-treated controls, with a corresponding increase in apoptotic cells. Early apoptotic cells (yellow-green fluorescence) were more prominent at 40 µM after 48 hours, while late apoptotic cells (orange-red nuclei) markedly increased at both concentrations and time points. Overall, RPG treatment induced an increased population of early apoptotic cells in all treatments in both cell lines (Fig. 2A, 2B). To corroborate these findings, apoptotic cell populations were quantitatively assessed using annexin V-FITC/PI dual staining followed by flow cytometry. As represented in Fig. 2C, MCF-7 cells treated with RPG showed an increase in early apoptotic populations from 0.25% in control to 8.07% at 40 µM and 16.19% at 80 µM. Similarly, A549 cells increased from 0.21% (control) to 7.71% and 9.18% at the respective doses (Fig. 2D). These flow cytometry results align with those of AO/EtBr, confirming the pro-apoptotic potential of RPG in both cell lines.

**Figure 2:**
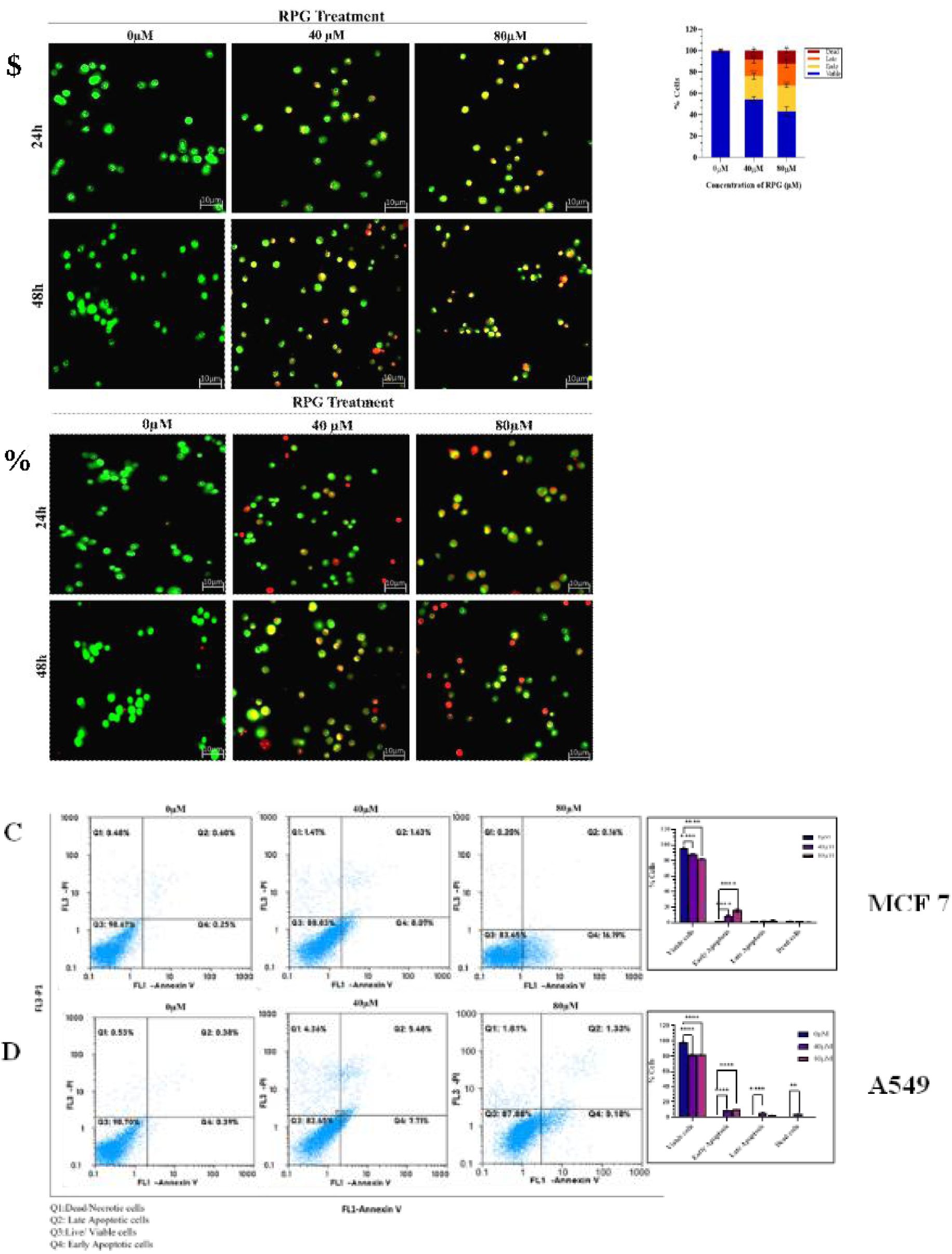
RPG-induced cellular-apoptotic changes in human breast and lung cancer cells: The cells were treated with two sub-lethal concentrations of RPG (40 μM and 80 μM). Figure A&B: AO/EtBr staining indicating the apoptotic changes induced by RPG in a time and concentration-dependent manner in MCF7 and A549 cells, respectively. C&D. Annexin V-FITC/PI double staining using flow cytometry, measuring apoptotic rate in RPG-treated MCF 7 AND A549, respectively. The graphs represent mean ± SD. Statistical significance is indicated as follows: *p < 0.05, **p < 0.01, ***p < 0.0001, and ****p < 0.00001, compared to the untreated control group.

### 3.3. RPG Induces DNA Damage and Promotes PARP Degradation in MCF-7 and A549 Cells

Considering the significant apoptotic response elicited by RPG in MCF-7 and A549 cancer cell lines, its ability to cause DNA damage was further assessed using the neutral comet assay. RPG treatment resulted in significant DNA damage in both cell types compared to vehicle-treated controls. This was characterized by a marked reduction in the percentage of head DNA with a corresponding increase in tail DNA content, indicative of DNA fragmentation. Moreover, a concentration-dependent increase in the olive tail moment was observed, further confirming RPG-induced genotoxic stress (Fig. 3A and B).

**Figure 3.**
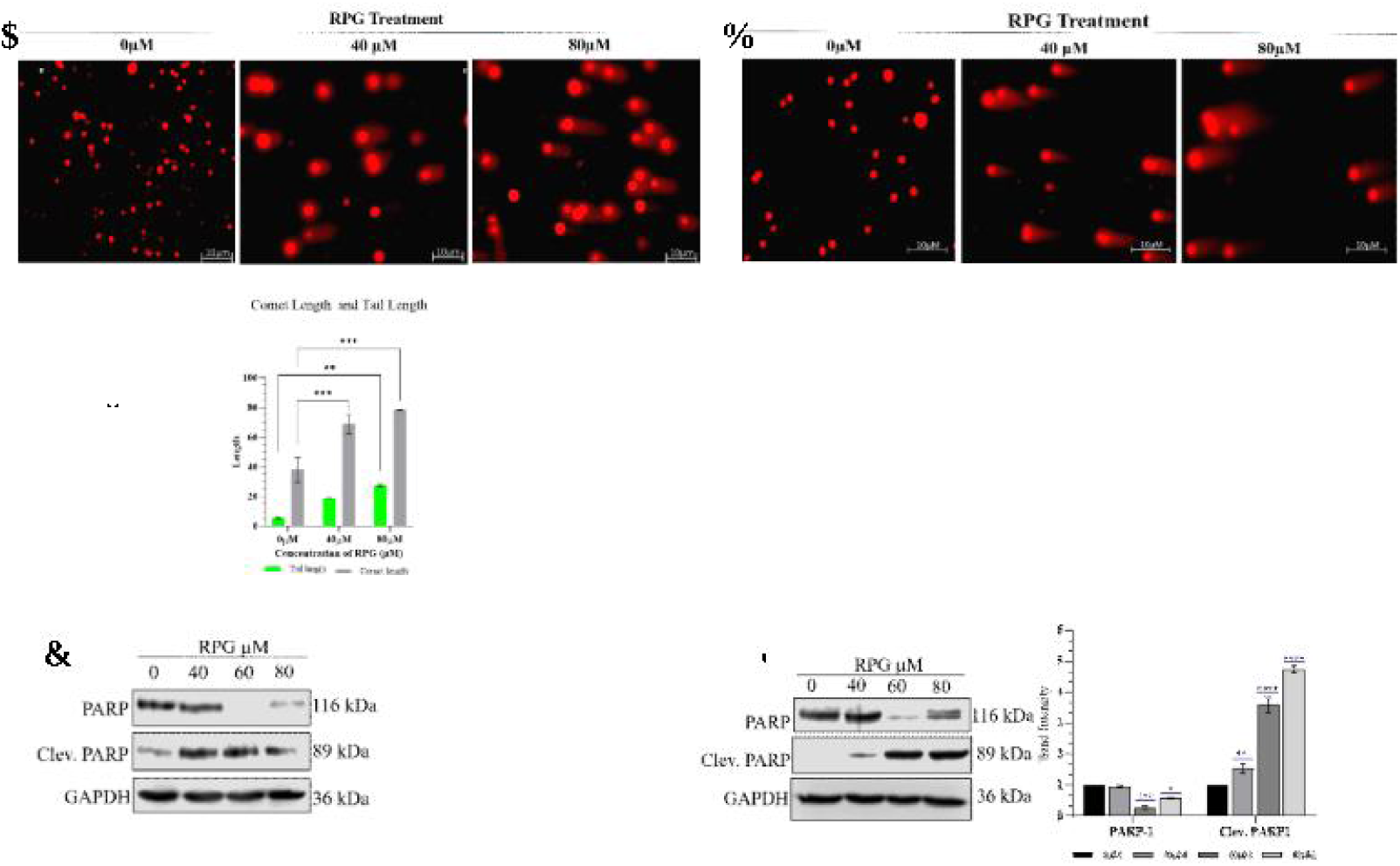
RPG-induced DNA damage is associated with altered PARP expression in MCF7 and A549 cell lines. Neutral comet assays were conducted on MCF7 and A549 cancer cells treated with two sublethal concentrations of RPG (40 and 80 µM) for 48 hours. Panels A and B illustrate representative comet images demonstrating DNA fragmentation in MCF7 and A549 cells post-treatment. Quantitative analyses show the percentage of head and tail DNA, tail length, comet length, tail moment, and Olive Tail Moment (OTM), based on 50 comets per condition. Panels C and D present Western blot analyses of PARP expression. RPG exposure led to notable changes in the levels of full-length and cleaved PARP, indicating DNA damage responses in both cell lines. Data are expressed as mean ± SD, with statistical analysis conducted using two-way ANOVA. Significant differences are indicated as **** p < 0.0001, *** p < 0.001, ** p < 0.01, and * p < 0.05, compared to the control group.

In order to further investigate the molecular response to RPG-induced DNA damage, western blot analysis was performed to assess the expression of poly (ADP-ribose) polymerase (PARP), a key marker of DNA damage and apoptosis. RPG treatment resulted in a concentration-dependent reduction in full-length PARP levels and a marked increase in cleaved PARP expression. (Fig. 3C & D). These results indicate that RPG induces significant DNA damage and promotes PARP cleavage, a critical event in the apoptotic pathway, in both MCF-7 and A549 cells.

### 3.4. RPG Triggers Apoptosis in MCF-7 and A549 Cells by Modulating Pro and Anti-apoptotic Proteins

The molecular mechanisms underlying RPG-induced apoptosis were examined by evaluating the expression levels of key pro- and anti-apoptotic proteins. Herein, MCF-7 and A549 cells were exposed to sublethal concentrations of RPG (40, 60, and 80 µM) for 48 hours, followed by Western blot analysis. As depicted in Fig. 4A and B, the immunoblotting results revealed a concentration-dependent increase in pro-apoptotic markers, including Bax and caspase-9, with a downregulation of the anti-apoptotic marker Bcl-2 in both cell lines. Simultaneously, a decrease in the expression of pro-caspase-3 was observed, accompanied by a corresponding increase in its active form, cleaved caspase-3, in RPG-treated A549 cells. In MCF-7 cells, RPG downregulated the expression of pro-caspase 7, indicating the downstream activation of the apoptotic pathway.

**Figure 4:**
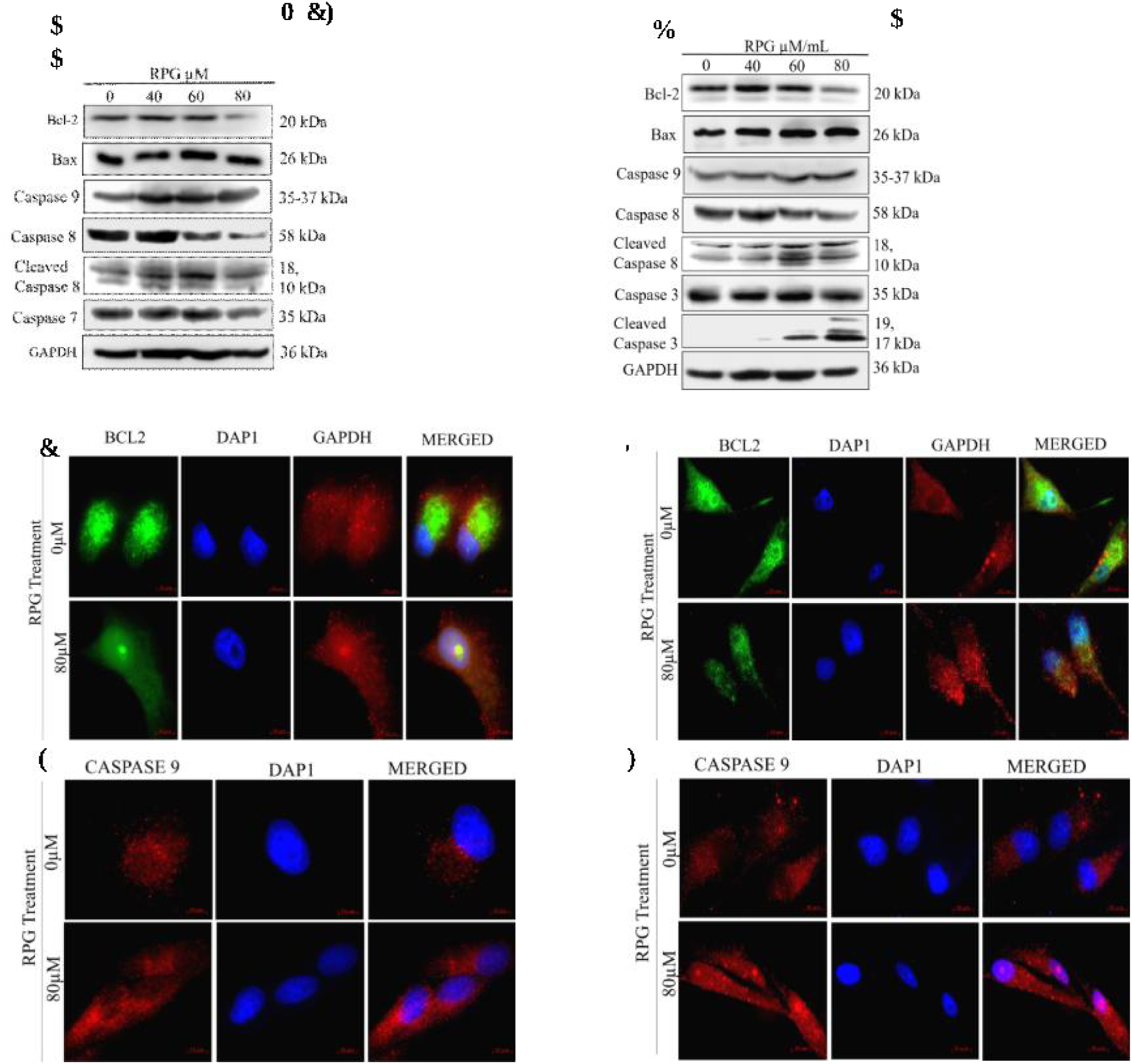
RPG mediates intrinsic and extrinsic apoptosis in MCF7 and A549 cells by altering the expression of pro- and anti-apoptotic proteins: Figure A and B: Western blot images depicting the expression levels of apoptotic markers in MCF7 and A549 cells following treatment with RPG at concentrations of 40, 60, and 80 μM. RPG treatment significantly downregulates the expression of BCL2 while upregulating the expression of pro-apoptotic proteins. (C&D) Immunofluorescence images showing the expression of Bcl-2 in RPG-treated MCF7 and A549 cells. (E, F) RPG-induced activation of the intrinsic apoptotic pathway is further supported by enhanced Caspase-9 expression, as demonstrated by immunofluorescence analysis. Bar diagrams represent the densitometric analysis of the corresponding western blot images, with GAPDH as the internal control. The bar represents means ± SD. Significance *p < 0.05, **p < 0.001 ***p < 0.0001

Additionally, the RPG treatment induced a downregulated expression of pro-caspase 8 with increased expression level of cleaved caspase-8 in both cell lines, indicating the activation of the extrinsic apoptotic pathway. To validate these findings, immunofluorescence staining was performed for Bcl-2 and caspase-9. In line with the western blot data, RPG-treated cells exhibited a pronounced decrease in Bcl-2 fluorescence intensity (Fig. 4C and 4D). In contrast, an enhanced caspase-9 expression (Fig. 4E and 4F) was observed in both MCF-7 and A549 cells. These results suggest that RPG effectively induces apoptosis in MCF-7 and A549 by activating intrinsic and extrinsic pathways.

### 3.5. RPG Impairs Colony Formation and Induces G1 Phase Arrest in MCF-7 and A549 cells

To evaluate the long-term antiproliferative effect of Repaglinide (RPG), a clonogenic assay was performed using sublethal concentrations (0–100 µM) in MCF-7 and A549 cells. As represented in Fig. 5A and B, RPG treatment led to a considerable, dose-dependent decrease in colony formation compared to vehicle-treated controls. A marked decline was observed at 40 µM and above concentrations, with near-complete inhibition of colony formation at the highest concentration tested. These findings are consistent with prior morphological assessments, further supporting the potent antiproliferative activity of RPG at higher concentrations in both cell lines.

**Figure 5:**
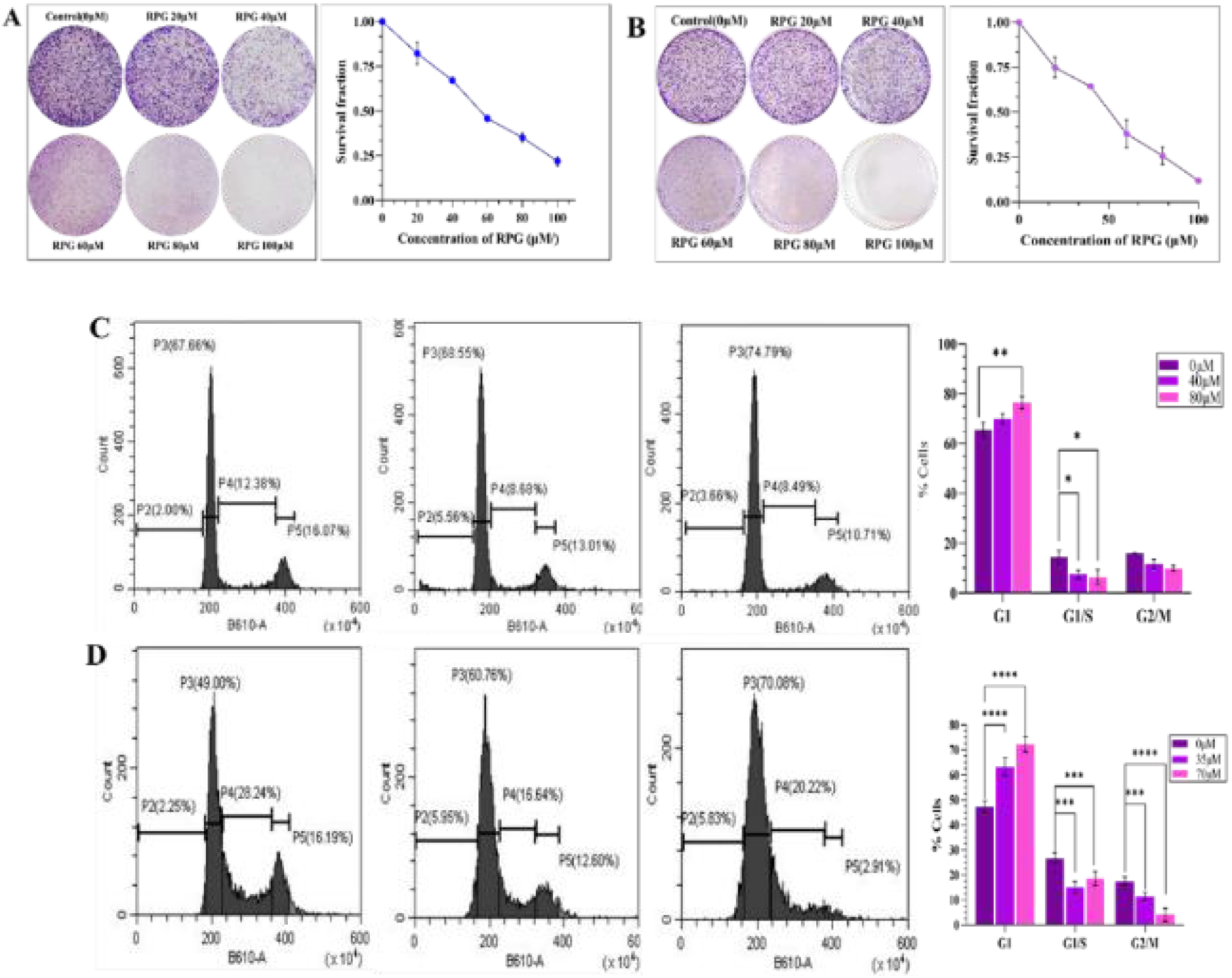
RPG inhibited colony formation and induced G1 cell cycle arrest in MCF7 and A549 cells: Figure A & B: Representative images from the colony formation assay showing MCF7 and A549 cells treated with increasing concentrations of RPG (0–100 µM). RPG treatment led to a concentration-dependent reduction in colony-forming ability in both cell lines. Figure C & D: Flow cytometric profiles illustrating the cell cycle distribution of MCF7 and A549 cells following treatment with RPG at 40 and 80 µM, respectively. RPG induced a marked G1 phase arrest in both cell types. The graph represents mean ± SD. The significance represented as *p < 0.05, **p < 0.001 ***p < 0.0001 and ****p< 0.00001

Furthermore, flow cytometric analysis was conducted to investigate the antiproliferative effect of RPG associated with cell cycle progression. RPG treatment led to G1 phase cell cycle arrest in both MCF-7 and A549 cells. In MCF-7 cells, the G1 phase population increased from 67.66% in control cells to 68.55% and 74.79% at 40 µM and 80 µM RPG, respectively, accompanied by a corresponding decrease in the G2/M phase (from 16.07% to 13.01% and 10.71%) (Fig. 5C). Similarly, in A549 cells, the G1 population rose from 49.00% in control cells to 60.76% and 70.08% following RPG treatment, with a concomitant decrease in G2/M phase from 16.19% to 12.60% and 2.91%, respectively (Fig. 5D). These findings indicate that RPG markedly inhibits colony-forming capacity by inducing G1 phase cell cycle arrest, thereby playing a key role in its antiproliferative activity in MCF-7 and A549 cells.

### 3.6. RPG Regulates the Expression of Cyclin-CDK Complexes and Modulates the PI3K/AKT/mTOR Pathway in Breast and Lung Cancer Cells

With the observed G1 phase arrest induced by Repaglinide (RPG) in MCF-7 and A549 cells, the expression of important cell cycle regulatory proteins was analyzed via western blotting. RPG treatment led to a marked, concentration-dependent downregulation of G1 phase cyclins and cyclin-dependent kinases, specifically cyclin D1 and CDK6, as well as cyclin E1 and CDK2, which are critical for G1/S phase transition. Additionally, the protein expressions of cyclin-dependent kinase inhibitors (CDKIs) and tumor suppressor, p53, were examined to elucidate the mechanism of RPG-induced cell cycle arrest. RPG treatment resulted in a significant upregulation of p53 and its downstream effector p21 in both cell lines, suggesting the involvement of the p53–p21 axis in enforcing G1 arrest. An increase in p16 expression was also noted, further contributing to cell cycle inhibition (Fig. 6A and B). The results from western blotting were further validated using immunofluorescence, where a notable decrease in fluorescent intensity of cyclin E and CDK2 was observed with a corresponding increase in the fluorescence intensity of p^21^ (Fig. 6C-H).

**Figure 6:**
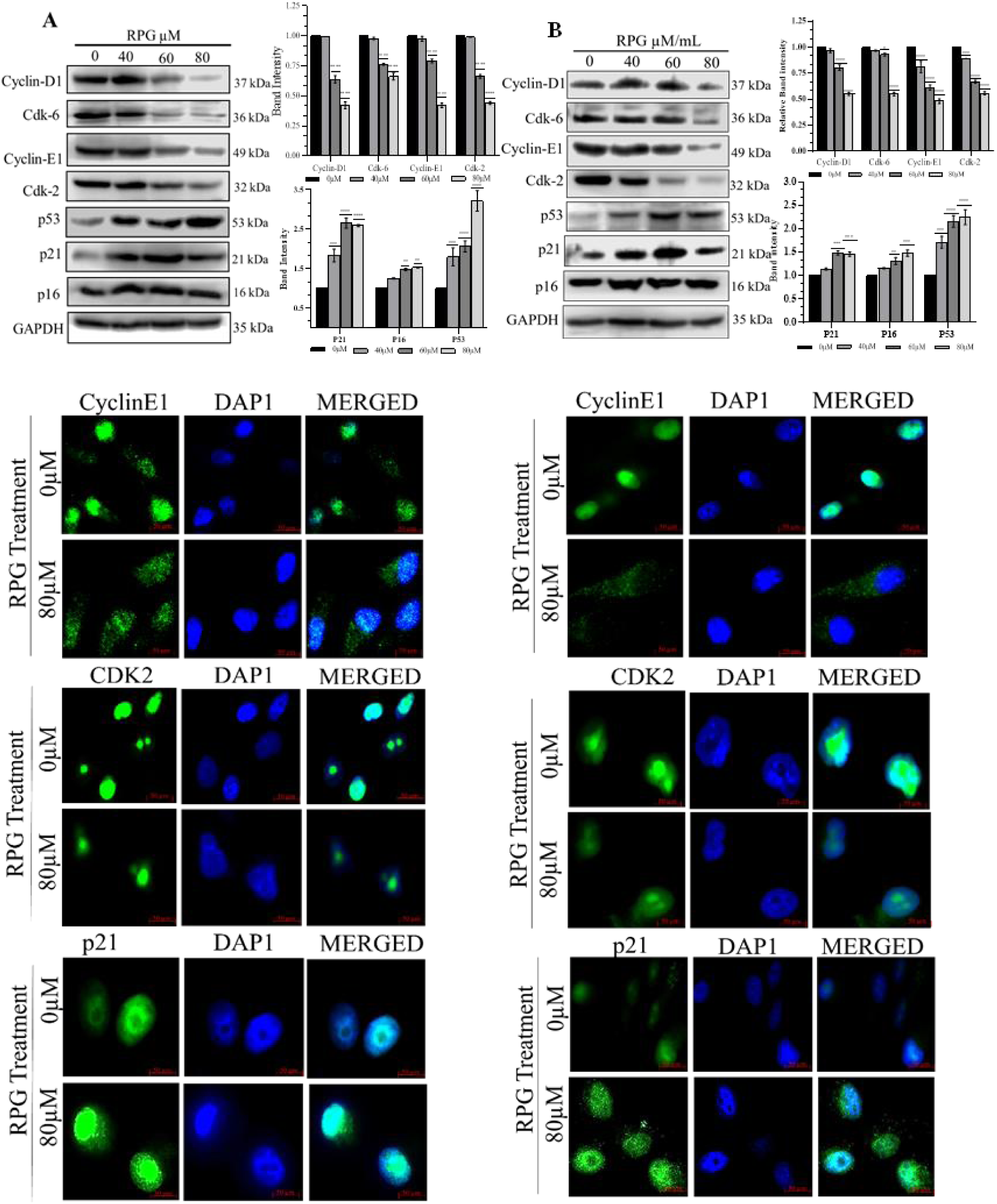
RPG treatment altered the expression of critical cell cycle regulatory proteins, leading to G1 phase arrest in both MCF7 and A549 cells. Cells exposed to three sublethal concentrations of RPG (40, 60, and 80 μM) were subsequently subjected to Western blotting. (A and B) Western blot images demonstrating a significant downregulation of Cyclin D1, Cyclin E1, CDK6, and CDK2, along with an upregulation of p53, p21, and p16 following RPG treatment. Immunofluorescence imaging further confirmed these findings, showing reduced expression of Cyclin E1 (C, D) and CDK2 (E, F), and increased expression of p21 (G, H) in both MCF7 and A549 cells. Data are presented as mean ± SD. Statistical significance is indicated as *p < 0.05, **p < 0.001, ***p < 0.0001, and ****p < 0.00001.

Given the pivotal role of the PI3K/AKT/mTOR signaling pathway in regulating cell proliferation and cell cycle progression, the expression of its key components was evaluated. RPG treatment resulted in a concentration-dependent downregulation of AKT, p-AKT, and mTOR, along with a concurrent upregulation of the tumor suppressor PTEN and p-PTEN, in both MCF-7 and A549 cells (Fig. 7A and B). Immunofluorescence studies further supported these findings, which confirmed reduced AKT and mTOR expression levels and enhanced PTEN expression in RPG-treated cells (Fig. 7C-H). Together, these results suggest that RPG induces G1 phase cell cycle arrest through a dual mechanism: downregulation of cyclin–CDK complexes via activation of the p53–p21 axis and inhibition of the PI3K/AKT/mTOR signaling pathway.

**Figure 7:**
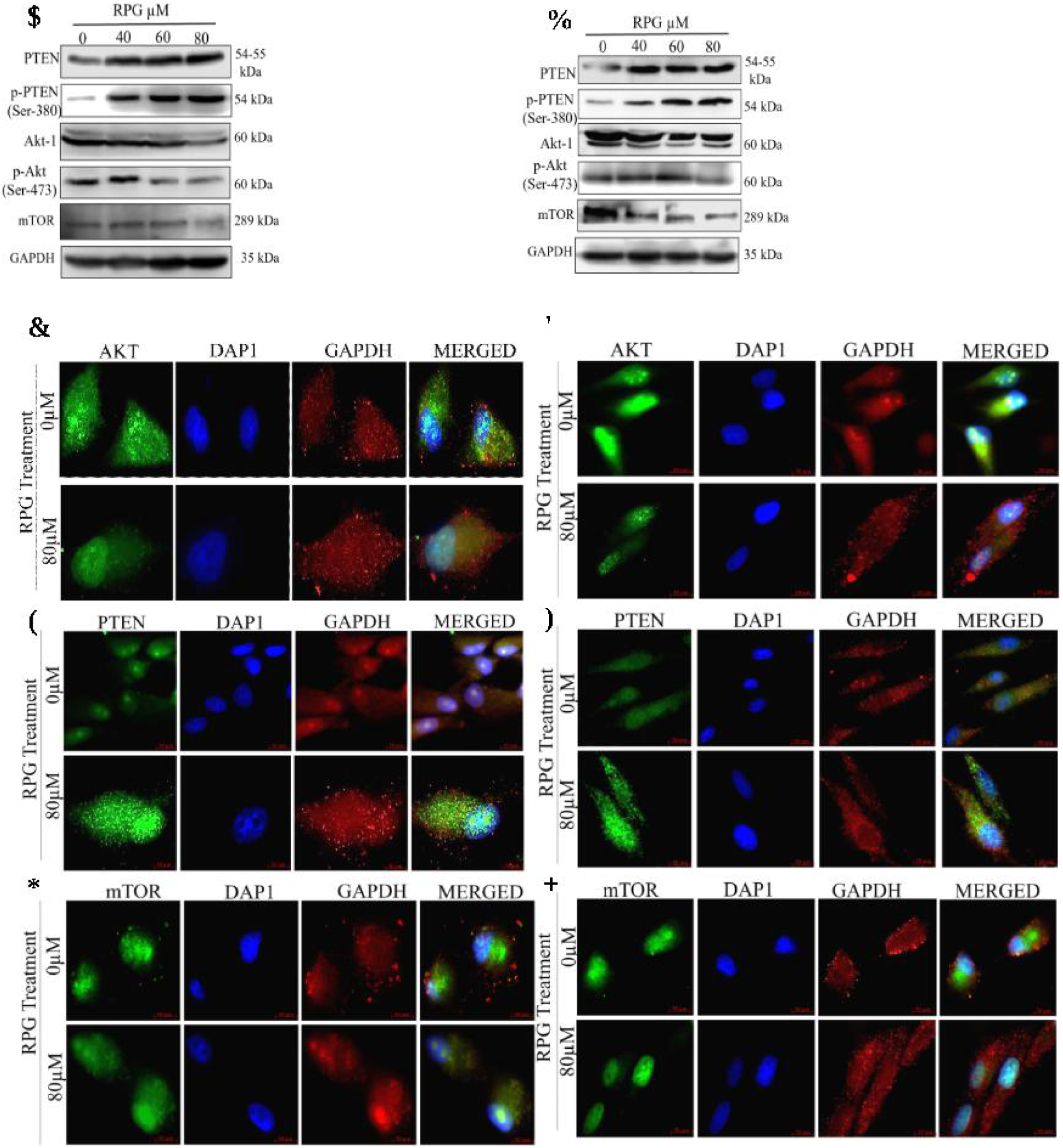
RPG modulates the expression of key proteins involved in cell proliferation in MCF7 and A549 cells. (A, B) Western blot analysis revealed an upregulation of PTEN and phosphorylated PTEN (p-PTEN), accompanied by a downregulation of AKT, phosphorylated AKT (p-AKT), and mTOR in RPG-treated MCF7 and A549 cells. (C, D) Immunofluorescence images confirmed increased PTEN expression in MCF7 and A549 cells, respectively. (E, F) Immunofluorescence analysis also showed reduced AKT expression following RPG treatment. (G, H) Similarly, mTOR expression was decreased in both cell lines as observed through immunofluorescence imaging. The graph represents mean ± SD. Data are presented as mean ± SD. Statistical significance is indicated as *p < 0.05, **p < 0.001, ***p < 0.0001, and ****p < 0.0000.

### 3.7. RPG promotes apoptosis and G1 phase arrest partially by regulating p53 expression

The involvement of p53 in RPG-mediated apoptosis and cell cycle arrest was further investigated using the specific p53 inhibitor pifithrin-α (PFR-α). MCF-7 and A549 cells were treated with RPG (80µM), PFR-α (10µM), or a combination of both, followed by protein expression analysis via Western blotting. Inhibition of p53 by PFR-α led to a significant upregulation of anti-apoptotic protein Bcl2 and a corresponding downregulation of the pro-apoptotic protein BAX. Moreover, p21 expression was markedly reduced in cells treated with PFR alone. Conversely, RPG treatment, both alone and in combination with PFR, suppressed BCL2 expression while enhancing the levels of BAX and p21; nonetheless, these modulations were less pronounced when RPG was combined with the p53 inhibitor. (Fig. 8A and B). These findings suggest that RPG may induce apoptosis and cell cycle arrest through mechanisms that are at least partially mediated by p53, but also involve p53-independent pathways.

**Figure 8:**
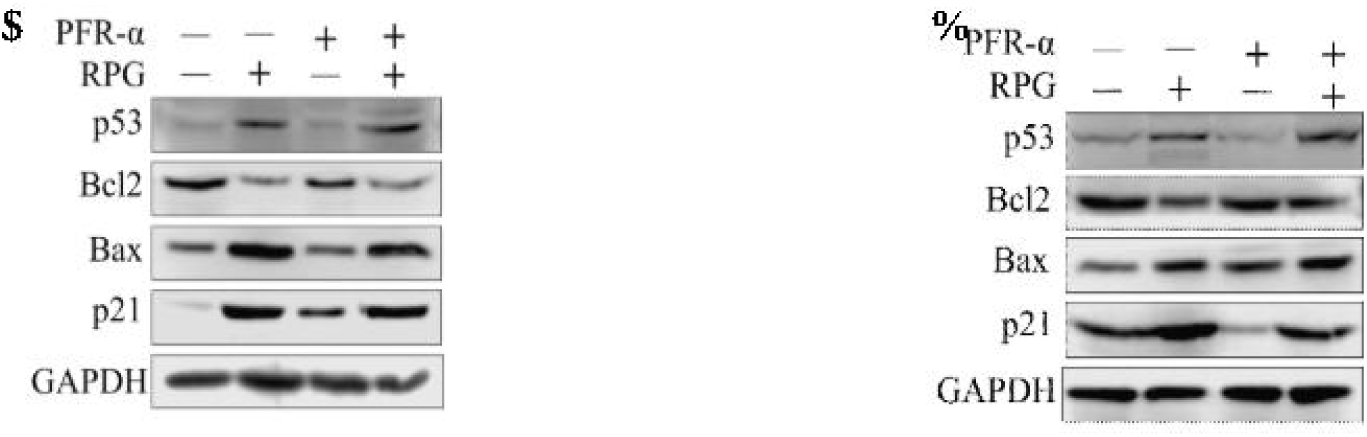
RPG Modulates Protein Expression Partially via the p53 Pathway. (A, B) MCF-7 and A549 cells were treated with a combination of Pifithrin-α (10 µM) and RPG (80 µM) for 48 hours. Protein levels of p53, Bcl-2, Bax, and p21 were evaluated by Western blot analysis. Quantitative bar graphs represent the densitometric analysis from three independent experiments, shown as mean ± SD. Statistical significance is indicated as *p < 0.05, **p < 0.01, ***p < 0.001, and ****p < 0.0001; * denotes within-group significance, while # indicates differences between treatment groups.

### 3.8. RPG Inhibits Migration of MCF-7 and A549 Cells

The anti-migratory effect of Repaglinide (RPG) was evaluated in MCF-7 and A549 cells using a scratch wound healing assay. Cells were treated with increasing sub-lethal concentrations of RPG (0–80 µM), and wound closure was assessed at 24 and 48 hours. RPG significantly reduced cell migration in both cell lines in a dose- and time-dependent manner. (Fig. 9A, B). At 24 hours, MCF-7 cells treated with 20 µM RPG exhibited 54% wound closure compared to 74% in controls, while A549 cells showed 70% closure versus 90% in controls. A further reduction in migration was observed at higher concentrations, with wound closure declining to 6% and 1% in MCF-7 and A549 cells, respectively, at 80 µM RPG. The decreased wound closure after 48h is mainly because of the cell death in the treated cells. After 48 hours, migration was nearly abolished at 60–80 µM RPG in both cell lines. Notably, A549 cells demonstrated a more pronounced sensitivity to RPG’s anti-migratory effects than MCF-7 cells. These findings are consistent with previous morphological and clonogenic assay results, reinforcing RPG’s potential to inhibit cancer cell migration.

**Figure 9:**
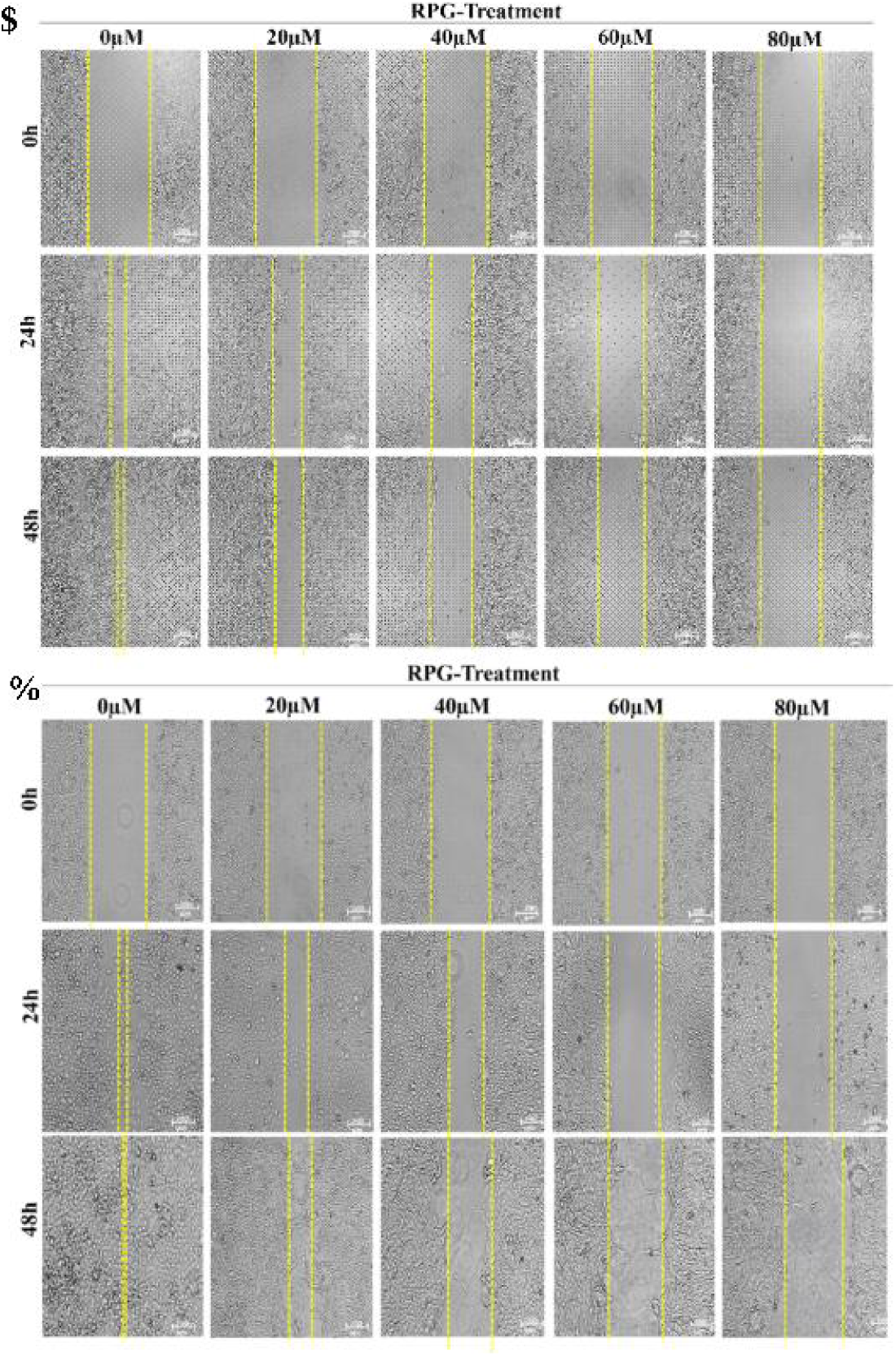
RPG inhibits cell migration as demonstrated by the scratch wound healing assay. (A) Representative images show wound closure in MCF7 cells treated with various concentrations of RPG at 24-, and 48-hours post-treatment. (B) Similar inhibitory effects on migration were observed in A549 cells. The corresponding graphs depict the percentage of wound area relative to RPG concentration at 24 and 48 hours. Statistical significance is indicated as *p < 0.05, **p < 0.01, ***p < 0.001, and ****p < 0.0001, where * denotes significance within groups and # indicates significance between groups.

### 3.9. Repaglinide Impairs Migration and Invasion to Reduce, potentially reduces cell metastasis in Breast and Lung Cancer Cells

Cell migration and invasion in MCF-7 and A549 cells after being subjected to RPG treatment (40 and 80 μM) were further validated using the transwell chamber assays. RPG treatment significantly reduced the cell migration and invasion in a concentration-dependent manner. In the migration assay, RPG reduced the number of migrating MCF-7 cells by 82.5% and 65% at 40 and 80 µM, respectively, while A549 cells showed reductions of 88% and 76% under the same conditions. (Fig. 10A) Similarly, in the invasion assay, RPG inhibited cell invasion through the extracellular matrix by 65% and 43% in MCF-7 cells and by 71% and 54% in A549 cells at 40 and 80 µM, respectively (Fig. 10 B). Substantiating the results of the scratch wound healing assay, these findings demonstrate that RPG significantly impairs cancer cell migration and invasion, underscoring its potential to suppress metastatic progression.

**Figure 10:**
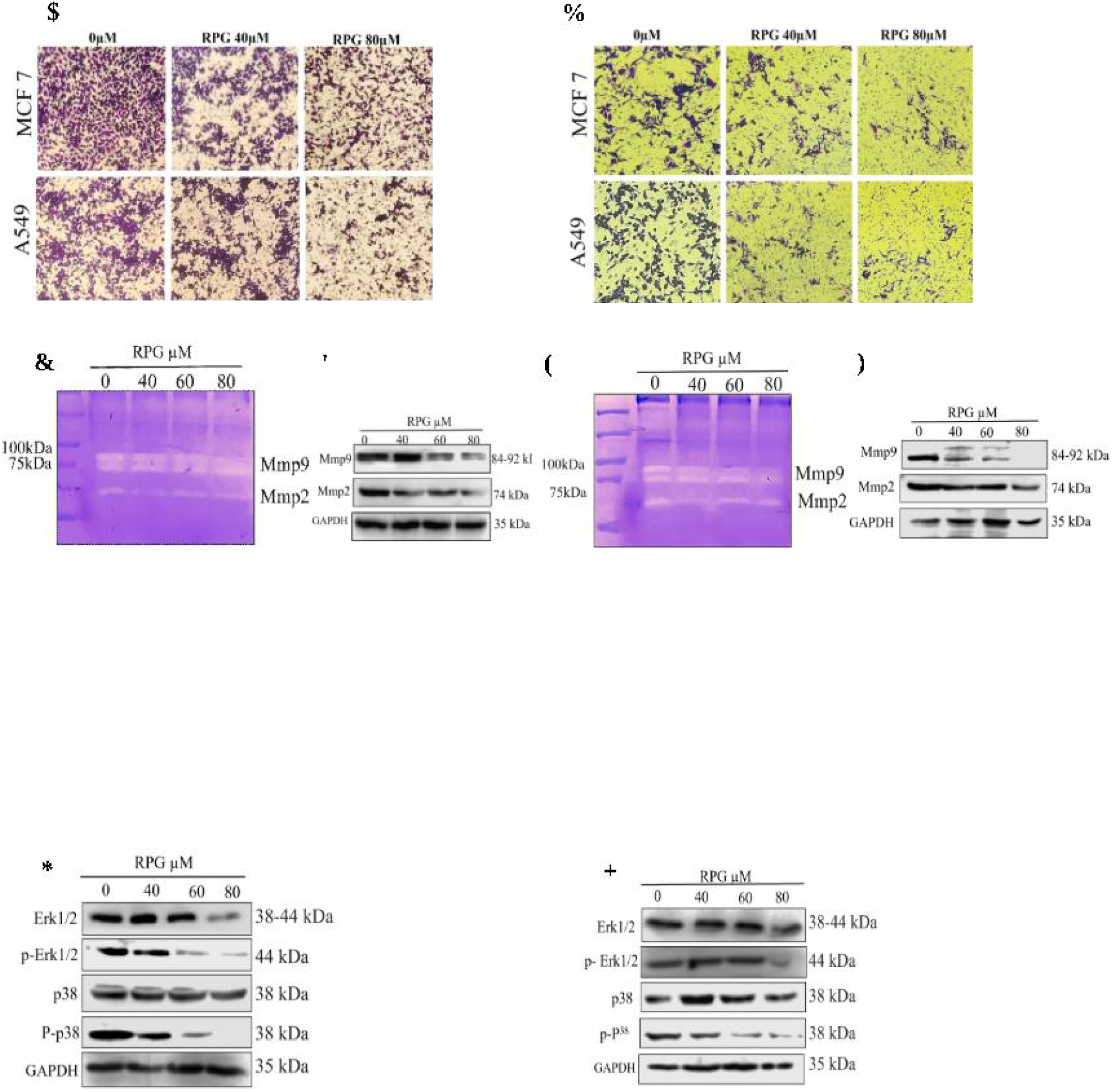
RPG inhibited cell migration via regulating the expressions and proteolytic activities of matrix metalloproteinases (MMP)-2 and MMP-9 in MCF7 and A549 cells: (A, B) Transwell migration and invasion assay demonstrated the anti-migratory effect of RPG on MCF7 and A549 cells. (C, E) Gelatin zymography showed a reduction in the proteolytic activity of MMP9 and MMP2 in MCF 7 and A549 cells following RPG treatment. (D, F) The protein expression levels of MMP-2 and MMP-9 in cells treated with three sub-lethal concentrations of RPG, with GAPDH serving as the loading control. (G, H) RPG treatment also altered the expression profiles of ERK and p38 MAPK, indicating potential involvement of these signaling pathways in RPG’s anti-migratory effects. The graph represents mean ±SD (n=3). Significant differences are indicated as **** p < 0.0001, *** p < 0.001, ** p < 0.01and * p < 0.05, compared to the control group.

**Figure 11:**
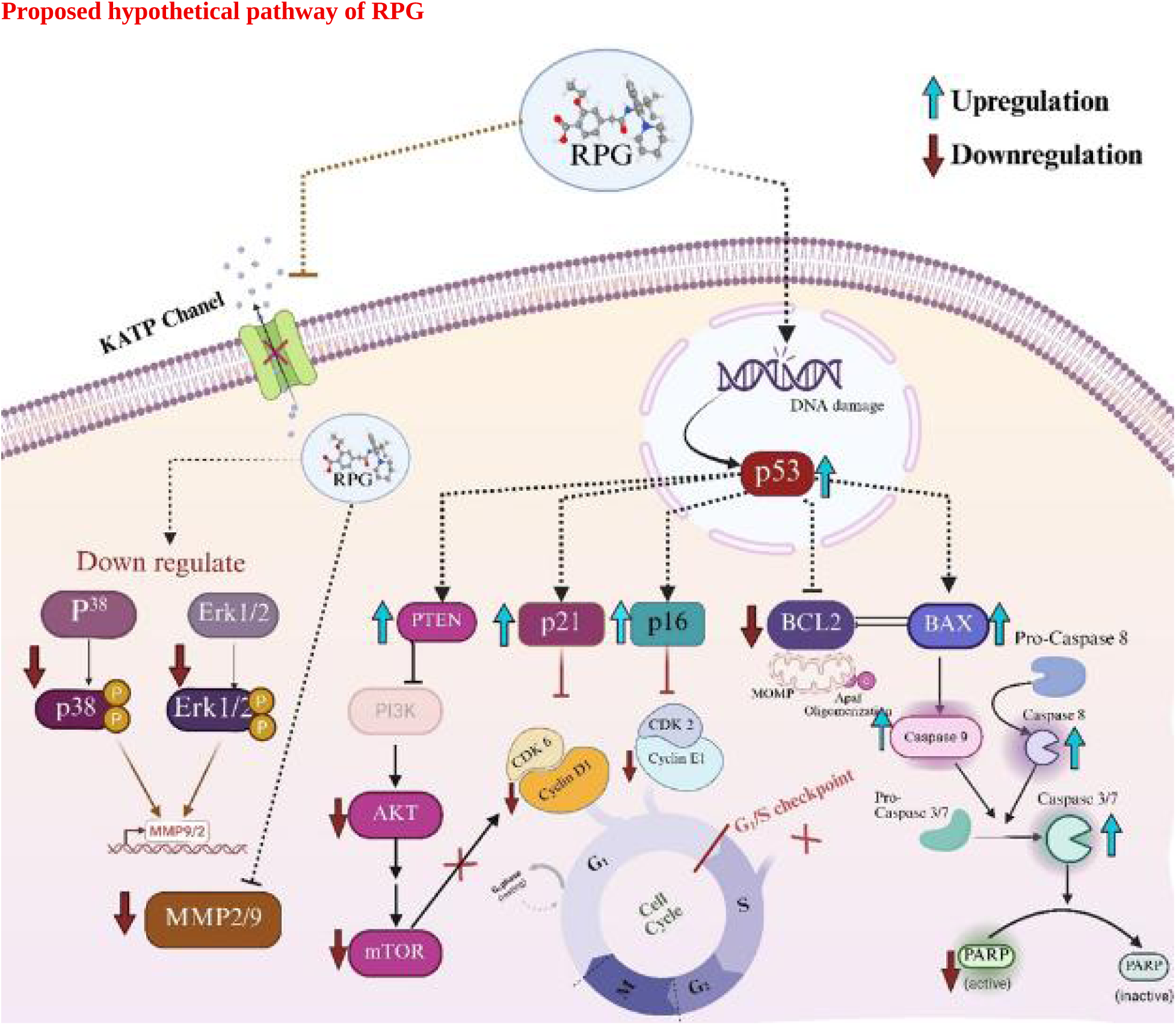
A schematic diagram of the hypothetical role of RPG in regulating various anticancer pathways in MCF7 and A549 cells. RPG- Repaglinide. (Created in BioRender. P k, H. (2025) https://BioRender.com/iv9ktx4)

### 3.10. Repaglinide Downregulates MMP-2 and MMP-9 Expression in MCF7 and A549 Cells

Given the key role of matrix metalloproteinases (MMPs) in cancer cell migration and invasion, the effect of RPG on MMP-2 and MMP-9 expression was evaluated using gelatin zymography and western blotting. RPG treatment led to a marked reduction in gelatinolytic activity, indicating decreased MMP activity. In A549 cells, MMP-2 and MMP-9 activity was reduced by approximately 60% and 40% at 60 and 80 µM RPG, respectively, with negligible change at 40 µM. (Fig. 10C) In MCF7 cells, all tested concentrations (40,60, and 80 µM) led to a moderate decrease in MMP activity, ranging between 60% and 70% (Fig. 10E). The effect of RPG on MMPs was further supported by western blot analysis, which confirmed a dose-dependent downregulation of MMP-2 and MMP-9 protein levels in both cell lines (Fig. 10D and F). These results suggest that RPG inhibits metastasis by modulating MMP expression in breast and lung cancer cells.

### 3.11. Repaglinide Modulates MAPK/ERK Signaling Pathway in a Cell-Specific Manner

To explore the role of the MAPK/ERK signaling pathway in RPG-mediated regulation of MMP expression, western blot analysis was performed on MCF-7 and A549 cells treated with increasing concentrations of RPG (40, 60, and 80 μM). In MCF-7 cells, RPG treatment resulted in a dose-dependent downregulation of total and phosphorylated forms of p38 and ERK proteins (Fig. 10.G). On the contrary, RPG treatment significantly downregulated the expression of total and phosphorylated p38, while ERK expression remained largely unaffected in A549 cells (Fig. 10.H). These results indicate that RPG modulates the MAPK/ERK signaling pathway in a cell-type-specific manner, potentially contributing to its inhibitory effects on MMP expression.

## 4. Discussion

Metabolic disorders and cancer share interconnected molecular and epidemiological foundations. Conditions such as Type 2 diabetes mellitus (T2DM), obesity, and metabolic syndrome are strongly associated with an increased risk of malignancies, including breast and lung cancers. Chronic hyperglycemia, hyperinsulinemia, insulin resistance, and systemic inflammation contribute to a pro-tumorigenic microenvironment that enhances proliferation and survival. Persistent activation of insulin and IGF signaling further stimulates key oncogenic pathways such as PI3K/Akt/mTOR and MAPK [6, 7,11]. These shared metabolic and cancer-associated mechanisms provide a strong rationale for investigating antidiabetic drugs as potential anticancer agents. In recent years, several glycemic regulators, including metformin and SGLT2 inhibitors, have demonstrated antiproliferative and pro-apoptotic effects across diverse tumor models, supporting the concept of anti-diabetic drug repurposing in oncology [14]. However, meglitinides such as repaglinide (RPG), despite their widespread clinical use as insulin secretagogues, remain comparatively underexplored in cancer research. Emerging evidence from glioblastoma and hepatocellular carcinoma models suggests that RPG may exert growth-inhibitory and pro-apoptotic effects independent of its glucose-lowering activity, highlighting the need to evaluate its broader anticancer potential carcinoma [18–20]. In the present study, we demonstrate that RPG significantly suppresses the proliferation of breast (MCF-7) and lung (A549) cancer cells through induction of p53-mediated G1 cell cycle arrest and activation of apoptotic signaling.

Assessment of cell viability is the primary step in evaluating the anticancer potential of any therapeutic candidate, as it provides a quantitative measure of growth inhibition and guides the selection of biologically relevant concentrations for downstream analyses [28]. Previous studies have reported that repaglinide (RPG) exhibits cytotoxic effects in certain malignancies, including glioblastoma (IC₅₀ ≈ 200 µM in L229 cells) and hepatocellular carcinoma (IC₅₀ ∼50 µM in HCCLM3 and MHCC97-H cells) 18,19]. However, its activity in breast and lung cancer models has not been clearly defined. In the present study, RPG markedly reduced the viability of MCF-7 and A549 cells in a dose- and time-dependent manner, with 48-hour IC₅₀ values of 100.8 ± 3.98 µM for MCF-7 and 104 ± 3 µM for A549 cell. These findings position RPG as an effective growth-inhibitory agent in epithelial cancers and highlight potential tumor-specific differences in drug sensitivity, possibly reflecting variations in metabolic state or apoptotic threshold. Importantly, the decline in cell viability was accompanied by distinct morphological alterations characteristic of apoptosis. Treated cells displayed shrinkage, loss of adherence, and membrane blebbing, particularly at concentrations ≥40 µM. Such changes are widely recognized as hallmarks of programmed cell death and have been consistently reported with established chemotherapeutic and pro-apoptotic agents in MCF-7 and A549 models [29,30].

Apoptotic evasion is a central hallmark of cancer progression and therapeutic resistance. Restoration of apoptotic signaling has therefore become a principal strategy in contemporary anticancer drug development [31]. In the present study, AO/EtBr staining and flow cytometric analyses using Annexin V-FITC/PI demonstrated that RPG-induced cytotoxicity was predominantly mediated through apoptosis rather than nonspecific necrosis. Previous studies in hepatocellular carcinoma have reported that RPG markedly downregulates Bcl-2 expression, thereby promoting apoptotic cell death [19]. Considering that Bcl-2 serves as a key anti-apoptotic regulator that maintains mitochondrial membrane stability and prevents cytochrome c release, its suppression represents a pivotal step in apoptotic initiation [32]. Consistent with this mechanistic framework, our findings indicate that RPG induces apoptosis through modulation of mitochondrial regulatory proteins, evidenced by a significant elevation in the Bax/Bcl-2 ratio in treated cells. Similar modulation of this apoptotic rheostat has been reported for other antidiabetic agents such as Metformin and Glibenclamide, [33,34], as well as several phytochemicals with established pro-apoptotic activity [35–37]. Furthermore, RPG-mediated activation of initiator caspase-9 along with downstream effector caspases (caspase-7 and cleaved caspase-3) confirms robust engagement of the intrinsic apoptotic pathway, while the concurrent activation of caspase-8 suggests additional involvement of extrinsic signaling. This coordinated activation of both apoptotic axes reflects a broad and potent pro-apoptotic mechanism. Such dual-pathway engagement is frequently associated with greater apoptotic efficiency and improved effectiveness in apoptosis-resistant malignancies [38]. The ability of RPG to trigger both apoptotic cascades demonstrates a comprehensive and potent mechanism that may significantly contribute to its anticancer efficacy.

The pro-apoptotic activity of RPG is strongly associated with the induction of DNA damage, which is a well-recognized upstream trigger of the intrinsic apoptotic pathway [39] (De Zio et al., 2013). DNA damage–associated apoptosis has been widely documented for metabolic agents such as Metformin and Canagliflozin across ovarian, colon, and hematological malignancies, where genomic instability precedes caspase activation and culminates in programmed cell death [40–42]. In agreement with these reports, RPG treatment resulted in enhanced DNA fragmentation and increased cleavage of PARP-1 in both MCF7 and A549 cells. Since PARP-1 cleavage is a well-established marker of execution-phase apoptosis and reflects irreversible commitment to cell death [43], these findings suggest that RPG-induced genomic damage may amplify intrinsic apoptotic signaling and promote efficient caspase activation. Comparable effects have been described for other meglitinides, such as Nateglinide, indicating that the class meglitinide may possess underrecognized genotoxic or DNA damage–sensitizing properties [44]. Notably, the ability of RPG to elicit DNA fragmentation and apoptotic markers at comparatively lower concentrations suggests enhanced potency relative to nateglinide. This concentration-dependent efficacy further highlights the potential of RPG as a promising candidate for repurposing in anticancer therapy.

Beyond the apoptotic effects, effective anticancer strategies increasingly focus on restricting tumor growth by interfering with cell cycle progression and attenuating key proliferative signaling networks [45]. In this context, the pronounced antiproliferative activity of RPG in MCF-7 and A549 cells underscores its potential as a multifunctional anticancer agent. The marked, dose-dependent reduction of colony-formation in the treated cells indicates that RPG impairs not only short-term viability but also the long-term replicative potential of cancer cells. This effect mirrors the growth-inhibitory outcomes reported for established chemotherapeutics such as doxorubicin in MCF-7 cells and paclitaxel in A549 cells [46–48], reinforcing the biological relevance of RPG-mediated growth suppression. Mechanistically, RPG induced a robust G1 phase arrest in both cell lines, accompanied by decreases in S and G2/M populations. This pattern parallels the cell cycle–modulatory effects described for metabolic agents such as Metformin and Canagliflozin, which similarly induce G0/G1 arrest across multiple cancer models [33,49,50]. G1 arrest is tightly regulated by the cyclin–cyclin-dependent kinase (CDK) axis, and RPG treatment resulted in substantial downregulation of cyclin D1, cyclin E1, CDK6, and CDK2, thereby disrupting both early and late G1 progression. Compared with metformin, which primarily targets cyclin D1 and CDK4/6 [33], RPG suppresses multiple cyclins and CDKs, indicating a coordinated and more comprehensive blockade of the G1 regulatory machinery. In parallel, RPG significantly upregulated cyclin-dependent kinase inhibitors (CKIs), including p21^Cip1/Waf1^, p16^INK4a^, and the tumor suppressor p53, indicating reinforcement of checkpoint control. The induction of p21 and p16 likely inhibits CDK2–cyclin E and CDK4/6 complexes, consolidating G1 arrest, while increased p53 expression suggests activation of upstream stress-responsive pathways that stabilize the checkpoint response. Tan et al. (2024) [19] similarly demonstrated RPG-mediated G1 arrest accompanied by activation of the p53–p21 pathway in hepatocellular carcinoma cells, supporting the reproducibility of this mechanism across distinct tumor types. The RPG-mediated cell cycle arrest was closely associated with attenuation of oncogenic proliferative signaling. In particular, RPG significantly influenced the PI3K/AKT/mTOR pathway, one of the most frequently dysregulated growth-regulatory cascades in cancer. In many cancers, the PI3K pathway and its downstream effectors, including AKT and mTOR, are frequently overactivated, whereas PTEN, a key negative regulator of this axis, is often downregulated [51]. In our study, RPG induced a dose-dependent reduction in AKT, p-AKT, and mTOR expression simultaneously upregulating PTEN and p-PTEN in both cell lines. Given that hyperactivation of AKT signaling promotes cyclin expression, CDK activation, and inhibition of p53-mediated checkpoint control [52], its attenuation by RPG provides a plausible upstream mechanism reinforcing G1 arrest and tumor suppressor activation. These findings parallel prior reports demonstrating that Metformin suppresses tumor growth via inhibition of PI3K/AKT/mTOR signaling in breast and other cancers [53]; however, the coordinated modulation of cyclins, CDKs, CKIs, p53, and PTEN observed here suggests that RPG exerts a more integrated regulatory influence over proliferative circuitry.

Within this interconnected framework, p53 emerges as a central molecular node orchestrating RPG-mediated anticancer responses in MCF-7 and A549 cells. As a master regulator of cellular stress signaling, p53 integrates pathways governing apoptosis, cell cycle arrest, DNA damage response, and metabolic adaptation [54]. RPG-induced upregulation of p53 was consistently associated with enhanced expression of downstream effectors, including p21 and BAX, alongside suppression of the anti-apoptotic protein Bcl-2, thereby functionally linking p53 activation to both G1 checkpoint regulation and mitochondrial apoptosis. The functional contribution of this axis was substantiated using the selective p53 inhibitor Pifithrin-α, [55] where partial reversal of RPG-induced apoptosis and cell cycle arrest confirmed a significant dependence on p53 transcriptional activity. Notably, the reported suppression of AKT signaling further supports mechanistic cross-talk, as reduced AKT activity can stabilize p53 through diminished MDM2-mediated degradation [56]. Thus, rather than acting through isolated molecular events, RPG appears to position p53 at the center of a coordinated regulatory network, integrating inhibition of proliferative signaling, reinforcement of G1 arrest, and activation of intrinsic apoptosis. Although partial persistence of these effects following p53 inhibition suggests auxiliary pathways may contribute, the collective molecular evidence supports p53 as the principal mediator through which RPG exerts its multifaceted anticancer activity.

In addition to its well-established anti-proliferative and pro-apoptotic effects, repaglinide (RPG) exhibited pronounced anti-migratory and anti-invasive activities, thereby expanding its anticancer repertoire. Consistent with earlier reports in glioblastoma and neuronal models [18,20] , as well as more recent findings in hepatocellular carcinoma [19], RPG significantly attenuated migratory capacity in MCF-7 and A549 cells, as demonstrated by wound healing and transwell invasion assays. These observations reinforce the notion that the anti-motility effects of RPG are not tumor-type specific but rather represent a broader biological property of the compound. Metastatic dissemination is critically dependent on dynamic cytoskeletal reorganization and proteolytic degradation of extracellular matrix (ECM) components, processes predominantly mediated by matrix metalloproteinases (MMPs), particularly MMP2 and MMP9 [57]. In agreement with this mechanistic framework, the present study demonstrates that RPG significantly suppresses both migration and invasion in breast and lung cancer cells. Importantly, our findings provide novel mechanistic insight by showing, for the first time, that RPG reduces both secreted and intracellular levels of MMP2 and MMP9, as evidenced by gelatin zymography and immunoblot analysis, thereby establishing a direct molecular link between RPG treatment and suppression of key ECM-remodeling enzymes. The regulation of MMP2 and MMP9 expression is tightly controlled by upstream signaling cascades, particularly the MAPK and PI3K/AKT pathways. Several natural and chemotherapeutic agents have been reported to inhibit tumor cell migration through modulation of these signaling networks. For example, genistein suppressed migration in pancreatic cancer cells via downregulation of MMP expression [58], and cyclophosphamide reduced invasiveness in breast cancer models through similar mechanisms [59]. Likewise, myricetin attenuated MMP-mediated migration by modulating ERK1/2 and p38 MAPK signaling [60], whereas apigenin exerted anti-invasive effects through inhibition of the PI3K/AKT pathway [61]. In line with these mechanistic paradigms, our study demonstrates that RPG markedly suppresses PI3K/AKT signaling and significantly reduces both total and phosphorylated forms of ERK1/2 and p38 MAPK. Given that transcriptional activation and enzymatic activity of MMP2 and MMP9 are governed by these upstream pathways, the concurrent inhibition of PI3K/AKT and MAPK cascades provides a coherent mechanistic explanation for the observed downregulation of MMP expression and the consequent impairment of invasive behavior.Collectively, when integrated with previous findings in glioblastoma, neuronal, and hepatocellular carcinoma models, the present study not only confirms the broad anti-migratory capacity of RPG but also advances current understanding by delineating a defined PI3K/AKT–MAPK–MMP2/9 signaling axis through which RPG exerts its anti-metastatic effects in breast and lung cancer cells

## Conclusion

The findings of the present study define a coordinated molecular mechanism by which repaglinide (RPG) suppresses tumor growth and progression in breast and lung cancer cells. Central to this regulatory network is the activation of p53, which integrates DNA damage signaling with mitochondrial apoptosis and G1 checkpoint control. RPG-induced DNA fragmentation and PARP-1 cleavage appear to act as upstream events that promote p53 stabilization, leading to transcriptional upregulation of p21 and Bax and thereby linking genomic stress to cell cycle arrest and intrinsic apoptotic execution. Simultaneously, inhibition of the PI3K/AKT/mTOR pathway, together with restoration of PTEN expression, attenuates proliferative and survival signaling. This suppression likely alleviates inhibitory constraints on p53 while limiting cyclin–CDK–mediated G1 progression, establishing a sustained antiproliferative state reflected in reduced clonogenic survival. Notably, the signaling pathways targeted by RPG also govern cytoskeletal dynamics and matrix remodeling. Reduced phosphorylation of AKT, ERK1/2, and p38 MAPK correlates with decreased MMP2 and MMP9 expression, providing mechanistic continuity between growth suppression and diminished migratory and invasive capacity. Overall, RPG does not appear to function through isolated molecular targets; rather, it orchestrates interconnected stress-responsive, proliferative, and metastatic signaling networks. This integrated modulation of apoptosis, checkpoint regulation, survival pathways, and invasion highlights the multifaceted anticancer potential of RPG and supports its further investigation as a repurposed therapeutic candidate in metabolically associated malignancies

## References

1. Tufail, M., Jiang, C.-H., & Li, N. (2024). Altered metabolism in cancer: insights into energy pathways and therapeutic targets. Molecular Cancer, 23(1). 10.1186/s12943-024-02119-3

2. Guneidy, R. A. (2026). Molecular basis of cancer chemoresistance: biochemical insights. Drug Metabolism Reviews, 1–27. 10.1080/03602532.2026.2613955

3. Anand, U., Dey, A., Chandel, A. K. S., Sanyal, R., Mishra, A., Pandey, D. K., De Falco, V., Upadhyay, A., Kandimalla, R., Chaudhary, A., Dhanjal, J. K., Dewanjee, S., Vallamkondu, J., & Pérez de la Lastra, J. M. (2022). Cancer chemotherapy and beyond: Current status, drug candidates, associated risks and progress in targeted therapeutics. Genes & Diseases, 10(4), 1367–1401. 10.1016/j.gendis.2022.02.007

4. Liu, B., Zhou, H., Tan, L., Siu, K. T. H., & Guan, X.-Y. (2024). Exploring treatment options in cancer: Tumor treatment strategies. Signal Transduction and Targeted Therapy, 9(1), 1–44. 10.1038/s41392-024-01856-7

5. Kulkarni, V. S., Alagarsamy, V., Solomon, V. R., Jose, P. A., & Murugesan, S. (2023). Drug Repurposing: An Effective Tool in Modern Drug Discovery. Russian Journal of Bioorganic Chemistry, 49(2), 157–166. 10.1134/s1068162023020139

6. Xia, Y., Sun, M., Huang, H., & Jin, W.-L. (2024). Drug repurposing for cancer therapy. Signal Transduction and Targeted Therapy, 9(1). 10.1038/s41392-024-01808-1

7. Dhas, Y., Biswas, N., M R, D., Jones, L. D., & Ashili, S. (2024). Repurposing metabolic regulators: antidiabetic drugs as anticancer agents. Molecular Biomedicine, 5(1), 40. 10.1186/s43556-024-00204-z

8. Rashmi, R., Singhal, R., & Misra, G. (2025). Diabetes driven effects on incidence and mortality of breast cancer: A meta-analysis. Public Health, 247, 105914. 10.1016/j.puhe.2025.105914

9. Sears, A. J., Wild, S. H., Mesa-Eguiagaray, I., Hall, P. S., & Figueroa, J. D. (2025). Breast cancer survival and mortality among women with type 2 diabetes: a retrospective cohort study. Scientific Reports, 15(1). 10.1038/s41598-025-08785-7

10. Azani, A., Avval, N. A., Meigoli, M. S. S., Imankhan, M., Asgari, P., Ebrahimisaraj, G., Morovatshoar, R., Hosseini, A. M., Mirzohreh, S. T., Nouri, M., Bostanabad, H. A., Behfar, Q., Bayrami, F., & Sharafi, M. (2025). Targeting the molecular crosstalk between diabetes and lung cancer for therapeutic intervention. Discover Oncology, 16(1). 10.1007/s12672-025-03264-x

11. Zhang, A. M. Y., Wellberg, E. A., Kopp, J. L., & Johnson, J. D. (2021). Hyperinsulinemia in Obesity, Inflammation, and Cancer. Diabetes & Metabolism Journal, 45(3). 10.4093/dmj.2020.0250

12. Sirtori, C. R., Castiglione, S., & Chiara Pavanello. (2024). METFORMIN: FROM DIABETES TO CANCER TO PROLONGATION OF LIFE. Pharmacological Research, 208, 107367–107367. 10.1016/j.phrs.2024.107367

13. Tilekar, K., Shelke, O., Upadhyay, N., Lavecchia, A., & Ramaa, C. S. (2022). Current status and future prospects of molecular hybrids with thiazolidinedione (TZD) scaffold in anticancer drug discovery. Journal of Molecular Structure, 1250, 131767. 10.1016/j.molstruc.2021.131767

14. Dutka, M., Bobiński, R., Francuz, T., Garczorz, W., Zimmer, K., Ilczak, T., Ćwiertnia, M., & Hajduga, M. B. (2022). SGLT-2 Inhibitors in Cancer Treatment—Mechanisms of Action and Emerging New Perspectives. Cancers, 14(23), 5811. 10.3390/cancers14235811

15. Chen, H., Zhao, L., Meng, Y., Qian, X., Fan, Y., Zhang, Q., Wang, C., Lin, F., Chen, B., Xu, L., Huang, W., Chen, J., & Wang, X. (2022). Sulfonylurea receptor 1-expressing cancer cells induce cancer-associated fibroblasts to promote non-small cell lung cancer progression. Cancer Letters, 536, 215611. 10.1016/j.canlet.2022.215611

16. Ray, S. D., Hussain, A., Aniqa Niha, Krmic, M., Jalshgari, A., Genis, D., & Jisha Reji. (2023). Anti-diabetic agents. Elsevier EBooks. 10.1016/b978-0-12-824315-2.01134-9

17. Tornio, A., Niemi, M., Neuvonen, P. J., & Backman, J. T. (2012). Drug interactions with oral antidiabetic agents: pharmacokinetic mechanisms and clinical implications. Trends in Pharmacological Sciences, 33(6), 312–322. 10.1016/j.tips.2012.03.001

18. Xiao, Z. X., Chen, R. Q., Hu, D. X., Xie, X. Q., Yu, S. B., & Chen, X. Q. (2017). Identification of repaglinide as a therapeutic drug for glioblastoma multiforme. Biochemical and Biophysical Research Communications, 488(1), 33–39. 10.1016/j.bbrc.2017.04.157

19. Tan, Y., Zhou, Y., Zhang, W., Wu, Z., Xu, Q., Wu, Q., Yang, J., Lv, T., Yan, L., Luo, H., Shi, Y., & Yang, J. (2024). Repaglinide restrains HCC development and progression by targeting FOXO3/lumican/p53 axis. Cellular Oncology, 47(4), 1167–1181. 10.1007/s13402-024-00919-9

20. Salcher, S., Spoden, G., Huber, J. M., Georg Golderer, Lindner, H., Ausserlechner, M. J., Kiechl-Kohlendorfer, U., Geiger, K., & Obexer, P. (2019). Repaglinide Silences the FOXO3/Lumican Axis and Represses the Associated Metastatic Potential of Neuronal Cancer Cells. Cells, 9(1), 1–1. 10.3390/cells9010001

21. Hernandez-Valencia, J., Garcia-Villa, E., Arenas-Hernandez, A., Garcia-Mena, J., Diaz-Chavez, J., & Gariglio, P. (2018). Induction of p53 Phosphorylation at Serine 20 by Resveratrol Is Required to Activate p53 Target Genes, Restoring Apoptosis in MCF-7 Cells Resistant to Cisplatin. Nutrients, 10(9), 1148. 10.3390/nu10091148

22. Bai, Z., Zhou, Y., Ye, X., Li, Y., Peng, Y., Guan, Q., Zhang, W., & Ma, L. (2021). Survivin suppression heightens BZML-induced mitotic catastrophe to overcome multidrug resistance by removing therapy-induced senescent A549/Taxol cells. Biochimica et Biophysica Acta (BBA) - Molecular Cell Research, 1869(2), 119174–119174. 10.1016/j.bbamcr.2021.119174

23. Riss, T. L., Moravec, R. A., Niles, A. L., Duellman, S., Benink, H. A., Worzella, T. J., & Minor, L. (2016, May 1). Cell Viability Assays. Nih.gov; Eli Lilly & Company and the National Center for Advancing Translational Sciences. https://www.ncbi.nlm.nih.gov/books/NBK144065/

24. Olive, P. L., & Banáth, J. P. (2006). The comet assay: a method to measure DNA damage in individual cells. Nature Protocols, 1(1), 23–29.10.1038/nprot.2006.5

25. Franken, N. A. P., Rodermond, H. M., Stap, J., Haveman, J., & van Bree, C. (2006). Clonogenic assay of cells in vitro. Nature Protocols, 1(5), 2315–2319. 10.1038/nprot.2006.339

26. Aroui, S., Aouey, B., Chtourou, Y., Meunier, A.-C., Fetoui, H., & Kenani, A. (2016). Naringin suppresses cell metastasis and the expression of matrix metalloproteinases (MMP-2 and MMP-9) via the inhibition of ERK-P38-JNK signaling pathway in human glioblastoma. Chemico-Biological Interactions, 244, 195–203. 10.1016/j.cbi.2015.12.011

27. Kielkopf, C. L., Bauer, W., & Urbatsch, I. L. (2020). Bradford Assay for Determining Protein Concentration. Cold Spring Harbor Protocols, 2020(4), pdb.prot102269. 10.1101/pdb.prot102269

28. Sánchez-Díez, M., Romero-Jiménez, P., Alegría-Aravena, N., Gavira-O’Neill, C. E., Vicente-García, E., Quiroz-Troncoso, J., González-Martos, R., Ramírez-Castillejo, C., & Pastor, J. M. (2025). Assessment of Cell Viability in Drug Therapy: IC50 and Other New Time-Independent Indices for Evaluating Chemotherapy Efficacy. Pharmaceutics, 17(2), 247. 10.3390/pharmaceutics17020247

29. Muhammed, Y., & Lazenby, R. A. (2024). Scanning ion conductance microscopy revealed cisplatin-induced morphological changes related to apoptosis in single adenocarcinoma cells. Analytical Methods, 16(4), 503–514. 10.1039/d3ay01827j

30. Cascione, M., De Matteis, V., Mandriota, G., Leporatti, S., & Rinaldi, R. (2019). Acute Cytotoxic Effects on Morphology and Mechanical Behavior in MCF-7 Induced by TiO2NPs Exposure. International Journal of Molecular Sciences, 20(14), 3594. 10.3390/ijms20143594

31. Kim, R., Kin, T., & Beck, W. T. (2024). Impact of Complex Apoptotic Signaling Pathways on Cancer Cell Sensitivity to Therapy. Cancers, 16(5), 984. 10.3390/cancers16050984

32. Qian, S., Wei, Z., Yang, W., Huang, J., Yang, Y., & Wang, J. (2022). The role of BCL-2 family proteins in regulating apoptosis and cancer therapy. Frontiers in Oncology, 12. 10.3389/fonc.2022.985363

33. Queiroz, E. A. I. F., Puukila, S., Eichler, R., Sampaio, S. C., Forsyth, H. L., Lees, S. J., Barbosa, A. M., Dekker, R. F. H., Fortes, Z. B., & Khaper, N. (2014). Metformin Induces Apoptosis and Cell Cycle Arrest Mediated by Oxidative Stress, AMPK and FOXO3a in MCF-7 Breast Cancer Cells. PLoS ONE, 9(5), e98207. 10.1371/journal.pone.0098207

34. Yan, B., Peng, Z., Xing, X., & Du, C. (2017). Glibenclamide induces apoptosis by activating reactive oxygen species dependent JNK pathway in hepatocellular carcinoma cells. Bioscience Reports, 37(5). 10.1042/bsr20170685

35. Xu, G., Yu, B., Wang, R., Jiang, J., Wen, F., & Shi, X. (2021). Astragalin flavonoid inhibits proliferation in human lung carcinoma cells mediated via induction of caspase-dependent intrinsic pathway, ROS production, cell migration and invasion inhibition and targeting JAK/STAT signalling pathway. Cellular and Molecular Biology, 67(2), 44–49. 10.14715/cmb/2021.67.2.7

36. Pandey, K., Tripathi, S. K., Panda, M., & Biswal, B. K. (2020). Prooxidative activity of plumbagin induces apoptosis in human pancreatic ductal adenocarcinoma cells via intrinsic apoptotic pathway. Toxicology in Vitro, 65, 104788. 10.1016/j.tiv.2020.104788

37. Faramarzi, F., Alimohammadi, M., Rahimi, A., Alizadeh-Navaei, R., Shakib, R. J., & Rafiei, A. (2022). Naringenin induces intrinsic and extrinsic apoptotic signaling pathways in cancer cells: A systematic review and meta-analysis of in vitro and in vivo data. Nutrition Research, 105, 33–52. 10.1016/j.nutres.2022.05.003

38. Wani, A. K., Akhtar, N., Mir, T. ul G., Singh, R., Jha, P. K., Mallik, S. K., Sinha, S., Tripathi, S. K., Jain, A., Jha, A., Devkota, H. P., & Prakash, A. (2023). Targeting Apoptotic Pathway of Cancer Cells with Phytochemicals and Plant-Based Nanomaterials. Biomolecules, 13(2), 194. 10.3390/biom13020194

39. De Zio, D., Cianfanelli, V., & Cecconi, F. (2013). New Insights into the Link Between DNA Damage and Apoptosis. Antioxidants & Redox Signaling, 19(6), 559–571. 10.1089/ars.2012.4938

40. Zhang, J., Zhou, P., Wu, T., Zhang, L., Kang, J., Liao, J., Jiang, D., Hu, Z., Han, Z., & Zhou, B. (2024). Metformin combined with cisplatin reduces anticancer activity via ATM/CHK2-dependent upregulation of Rad51 pathway in ovarian cancer. Neoplasia, 57, 101037. 10.1016/j.neo.2024.101037

41. Wang, S., Sui, M., Chen, Q., Guo, J., Yang, H., Zhou, Y., Ji, M., Cheng, Y., & Hou, P. (2024). Engineering PD-L1 targeted liposomal canagliflozin achieves multimodal synergistic cancer therapy. Chemical Engineering Journal, 498, 155074. 10.1016/j.cej.2024.155074

42. Song, Y., Chen, S., Xiang, W., Xiao, M., & Xiao, H. (2021). The mechanism of treatment of multiple myeloma with metformin by way of metabolism. Archives of Medical Science, 17(4), 1056–1063. 10.5114/aoms.2020.101305 40

43. Ray Chaudhuri, A., & Nussenzweig, A. (2017). The multifaceted roles of PARP1 in DNA repair and chromatin remodelling. Nature Reviews Molecular Cell Biology, 18(10), 610–621. 10.1038/nrm.2017.53

44. Samet z, G ekerci, Furkan ksel, & Tekin, S. (2023). Cytotoxic and genotoxic effects of nateglinide on human ovarian, prostate, and colon cancer cell lines. Annals of Medical Research, 0, 1–1. 10.5455/annalsmedres.2023.02.062

45. Suski, J. M., Braun, M., Strmiska, V., & Sicinski, P. (2021). Targeting cell-cycle machinery in cancer. Cancer Cell, 39(6), 759–778. 10.1016/j.ccell.2021.03.010

46. Xu, J., Zhang, Z., Hu, H., Yang, Y., Xiao, C., Xi, L., Lu, J., Tian, S., & Zhao, H. (2024). Synergistic antitumor effects of Peiminine and Doxorubicin on breast cancer through enhancing DNA damage via ZEB1. Biomedicine & Pharmacotherapy, 173, 116353. 10.1016/j.biopha.2024.116353

47. Effat, H., Abosharaf, H. A., & Radwan, A. M. (2024). Combined effects of naringin and doxorubicin on the JAK/STAT signaling pathway reduce the development and spread of breast cancer cells. Scientific Reports, 14(1), 2824. 10.1038/s41598-024-53320-9

48. Li, X.-Q., Ren, J., Wang, Y., Su, J.-Y., Zhu, Y.-M., Chen, C.-G., Long, W.-G., Jiang, Q., & Li, J. (2020). Synergistic killing effect of paclitaxel and honokiol in non-small cell lung cancer cells through paraptosis induction. Cellular Oncology, 44(1), 135–150. 10.1007/s13402-020-00557-x

49. Yamamoto, L., Yamashita, S., Nomiyama, T., Kawanami, T., Hamaguchi, Y., Shigeoka, T., Horikawa, T., Tanaka, Y., Yanase, T., Kawanami, D., & Iwasaki, A. (2021). Sodium-glucose cotransporter 2 inhibitor canagliflozin attenuates lung cancer cell proliferation in vitro. Diabetology International, 12(4), 389–398. 10.1007/s13340-021-00494-6

50. Shameem, M., Alireza Jian Bagherpoor, Nakhi, A., Dosa, P., Georg, G., & Fekadu Kassie. (2023). Mitochondria-targeted metformin (mitomet) inhibits lung cancer in cellular models and in mice by enhancing the generation of reactive oxygen species. Molecular Carcinogenesis, 62(11), 1619–1629. 10.1002/mc.23603

51. Peng, Y., Wang, Y., Zhou, C., Mei, W., & Zeng, C. (2022). PI3K/Akt/mTOR Pathway and Its Role in Cancer Therapeutics: Are We Making Headway? Frontiers in Oncology, 12. 10.3389/fonc.2022.819128

52. Glaviano, A., Song, A., Julia, Y., Kenneth Chun-Yong Yap, Jacot, W., Jones, R. H., Eng, H., Nair, M. G., Pooyan Makvandi, Geoerger, B., Kulke, M. H., Baird, R. D., Prabhu, J. S., Carbone, D., Pecoraro, C., Boon, D., Sethi, G., Cavalieri, V., Lin, K., & Javidi-Sharifi, N. (2023). PI3K/AKT/mTOR signaling transduction pathway and targeted therapies in cancer. Molecular Cancer, 22(1). 10.1186/s12943-023-01827-6

53. Alimova, I. N., Liu, B., Fan, Z., Edgerton, S. M., Dillon, T., Lind, S. E., & Thor, A. D. (2009). Metformin inhibits breast cancer cell growth, colony formation and induces cell cycle arrest in vitro. Cell Cycle, 8(6), 909–915. 10.4161/cc.8.6.7933

54. Hernández Borrero, L. J., & El-Deiry, W. S. (2021). Tumor suppressor p53: Biology, signaling pathways, and therapeutic targeting. Biochimica et Biophysica Acta (BBA) - Reviews on Cancer, 1876(1), 188556. 10.1016/j.bbcan.2021.188556

55. Zhu, J., Singh, M., Selivanova, G., & Peuget, S. (2020). Pifithrin-α alters p53 post-translational modifications pattern and differentially inhibits p53 target genes. Scientific Reports, 10(1). 10.1038/s41598-020-58051-1

56. Chibaya, L., Karim, B., Zhang, H., & Jones, S. N. (2021). Mdm2 phosphorylation by Akt regulates the p53 response to oxidative stress to promote cell proliferation and tumorigenesis. Proceedings of the National Academy of Sciences, 118(4). 10.1073/pnas.2003193118

57. Mustafa, S., Koran, S., & AlOmair, L. (2022). Insights Into the Role of Matrix Metalloproteinases in Cancer and its Various Therapeutic Aspects: A Review. Frontiers in Molecular Biosciences, 9. 10.3389/fmolb.2022.896099

58. Bi, Y., Min, M., Shen, W., & Liu, Y. (2018). Genistein induced anticancer effects on pancreatic cancer cell lines involves mitochondrial apoptosis, G 0 /G 1 cell cycle arrest and regulation of STAT3 signalling pathway. Phytomedicine, 39, 10–16. 10.1016/j.phymed.2017.12.001

59. Izdebska, M., Zielińska, W., Krajewski, A., Hałas-Wiśniewska, M., Mikołajczyk, K., Gagat, M., & Grzanka, A. (2021). Downregulation of MMP-9 Enhances the Anti-Migratory Effect of Cyclophosphamide in MDA-MB-231 and MCF-7 Breast Cancer Cell Lines. International Journal of Molecular Sciences, 22(23), 12783. 10.3390/ijms222312783

60. Kang, H. R., Moon, J. Y., Ediriweera, M. K., Song, Y. W., Cho, M., Kasiviswanathan, D., & Cho, S. K. (2020). Dietary flavonoid myricetin inhibits invasion and migration of radioresistant lung cancer cells (A549-IR) by suppressing MMP-2 and MMP-9 expressions through inhibition of the FAK-ERK signaling pathway. Food Science & Nutrition, 8(4), 2059–2067.10.1002/fsn3.1495

61. Zhou, Z., Tang, M., Liu, Y., Zhang, Z., Lu, R., & Lu, J. (2017). Apigenin inhibits cell proliferation, migration, and invasion by targeting Akt in the A549 human lung cancer cell line. Anti-Cancer Drugs, 28(4), 446–456. 10.1097/cad.0000000000000479

62. Created in BioRender. P k, H. (2025) https://BioRender.com/iv9ktx4

